# Yeast aging from a dynamic systems perspective: Analysis of single cell trajectories reveals significant interplay between nuclear size scaling, proteasome dynamics, and mitochondrial morphology

**DOI:** 10.1101/2025.03.11.642143

**Authors:** Michael Mobaraki, Changhui Deng, Jiashun Zheng, Hao Li

## Abstract

Yeast replicative aging is cell autonomous and thus a good model for mechanistic study from a dynamic systems perspective. Utilizing an engineered strain of yeast with a switchable genetic program to arrest daughter cells (without affecting mother cell divisions) and a high throughput microfluidic device, we systematically analyze the dynamic trajectories of thousands of single yeast mother cells throughout their lifespan, using fluorescent reporters that cover a range of biological processes, including some major aging hallmarks. We found that the markers of proteostasis stand out as most predictive of the lifespan of individual cells. In particular, nuclear proteasome concentration at middle age is a good predictor. We found that cell size (measured by area) grows linearly with time, and that nuclear size grows in proportion to maintain isometric scaling in young cells. As the cells become older, their nuclear size increases faster than linear and isometric size scaling breaks down. We observed that proteasome concentration in the nucleus exhibits dynamics very different from that in cytoplasm, with much more rapid decrease during aging; such dynamic behavior can be accounted for by the change of nuclear size in a simple mathematical model of transport. We hypothesize that the gradual increase of cell size and the associated nuclear size increase lead to the dilution of important nuclear factors (such as proteasome) that drives aging. We also show that perturbing proteasome changes mitochondria morphology and function, but not vice versa, potentially placing the change of proteosome upstream of the change of mitochondrial phenotypes. Our study produced large scale single cell dynamic data that can serve as a valuable resource for the aging research community to analyze the dynamics of other markers and potential causal relations between them. It is also a useful resource for building and testing physics/AI based models that identify early dynamics events predictive of lifespan and can be targets for longevity interventions.

## Introduction

Budding yeast has been a canonical model for cellular aging study, owing to its short lifespan, genetic tractability, and conservation of biological pathways across species^1–5^. In addition, during replicative aging, yeast cells manifest many hallmarks of aging observed in multi-cellular organism^2^. Application of molecular genetic to this model systems has led to the discovery of many genes whose perturbation significantly extends lifespan, leading to important insight into the biology of aging^3,6,7^. However, it is still very challenging to piece together the genetic evidences to arrive at a wholistic picture of how cells age and die, as aging is a complex and multifactorial process.

With the invention of microfluidic devices for single cell aging study in yeast, it has become possible to analyze the process of aging from a dynamic systems perspective^8–12^. Yeast is particularly amenable for such analysis as replicative aging is cell autonomous; thus, each single cell can be treated as a separate dynamic system with a constant environment (maintained in microfluidic devices), circumventing the complexity of cell-cell interaction in multi-cellular organisms.

With single cell tracking and molecular reporters, it is possible to observe how various cellular parameters change over time throughout the lifespan of the yeast mother cells^8–10,13–15^. Examples of observables include cell and organelle size and shape, cell cycle time, and molecular readout of various cellular processes such as proteostasis, genome stability, and mitochondria function. Such observations allows the tracking of the “dynamic state” of a cell over time throughout its lifespan, with the potential to identify early dynamic events that might be causal to the eventual loss of cellular homeostasis and cellular aging phenotypes. Several recent studies on yeast aging took this approach and discovered interesting properties of dynamic trajectories that determine the terminal cell fate. Such trajectories can be altered with genetic/environmental perturbations, leading to significantly lifespan extension^9,10,13,16,17^.

Here we systematically analyze the dynamic trajectory of thousands of single yeast mother cells throughout their lifespan, using fluorescent reporters that cover a range of biological processes, including the majority of aging hallmarks, such as genomic instability, loss of proteostasis, mitochondrial dysfunction, epigenetic alteration, de-regulated nutrient sensing, and altered stress response . We have constructed 34 single reporter strains, and 29 double reporter strains, 15 knockouts, and 20 inducible perturbations to analyze the dynamics of different processes and the interrelations between them.

Analysis of these single cell trajectories revealed several interesting phenomena that can be used to construct a hypothesis on how the gradual drift of the cellular state leads to the eventual loss of homeostasis and cellular aging. By searching for markers for lifespan that change with age and is predictive of the lifespan of individual cells, we found that reporters for proteostasis stand out among 34 reporters we analyzed. In particular, nuclear proteosome concentration at mid age is a good predictor of lifespan. We observed that the cell size (measured by the surface area) grows linearly with time, and that initially nuclear size (area) also grows linearly to maintain isometric size scaling. However, such scaling breaks down as cells become old, with nuclear size increasing faster than linear. We found that proteasome concentration in the nucleus exhibits dynamics very different from that in cytoplasm, with much more rapid decrease during aging; such dynamic behavior can be accounted for by the change of nuclear size in a simple mathematical model of nuclear transport. Time dependent over-expression of proteasome leads to significant increase of nuclear size, pointing to potential negative feedback to maintain nuclear size/nuclear proteasome homeostasis. Thus, our dynamic trajectory analysis revealed interesting interplay between size scaling, nuclear size dynamics, and the dynamics of nuclear proteasome. We hypothesize that the gradual increase of cell size and the associated nuclear size leads to the dilution of important nuclear factors (such as proteasome) that drives aging, a hypothesis that needs to be further tested.

Our study generated a large-scale, single-cell dynamic dataset, providing a valuable resource for the aging research community to analyze the temporal dynamics of various cellular markers and infer potential causal relationships between them. For instance, we found that proteasome perturbation significantly alters mitochondrial morphology, whereas mitochondrial morphological changes do not reciprocally impact the proteasome, suggesting a directional causal relationship between these two hallmarks of aging. In addition to serving as a foundation for AI-driven classification and segmentation, our dataset enabled the development of high-throughput AI models capable of identifying and categorizing distinct cellular topologies. Furthermore, our single cell observations maybe used to derive physics- and AI-based models to detect early dynamic features predictive of lifespan, which may serve as promising targets for longevity interventions.

## Results

### Tracking the dynamics of single yeast cells aging using a high-throughput imaging platform

To perform the high-throughput single cell imaging, we used an engineered strain with a daughter arresting program (DAP), previously developed in the Li lab^18^. Upon switching media, DAP is turned on and stops daughter cells from budding while leaving mother cell budding unaffected. This is achieved through a glucose repressible promoter that drives an essential and long-lived membrane protein. Upon switching to glucose media, the expression of the protein is repressed but mother cells already have the protein in its cell membrane. Because daughter cells do not inherit membrane from its mother (they synthesize new membranes), they do not have this protein in their membrane and their growth is arrested. Using DAP strain, we were able to track mother cells without the need of removing daughter cells, which drastically simplifies time lapsed imaging in microfluidic devices (no flow and no pumps needed, and no clogging issues).

To perform high throughput time-lapsed imaging, we combined the DAP strain with a high-throughput microfluidic device previously developed in the Li lab^18^. This device is made of a 96-wells plate interfacing with a microfluidic device with a two dimensional array of 32 modules, each aligned to three wells: an inlet and an outlet each connecting to a well, and an observational area (microfluidic channels and micro-structures) aligned to the middle well in between (Figure 1A,B). There are 220 microstructures in each module to trap single mother cells (Figure 1B). This device combines the advantage of 96 wells plate for cell loading and liquid handling and microfluidic device’s ability for single cell tracking. At its maximum capacity, this device can be used to analyze 32 different strains/conditions in parallel, each allowing the tracking of ∼200 cells. We have experimented with various cell loading protocols to optimize the process so that on average ∼80% of the microstructures can be loaded with one single mother cell.

**Fig 1.**
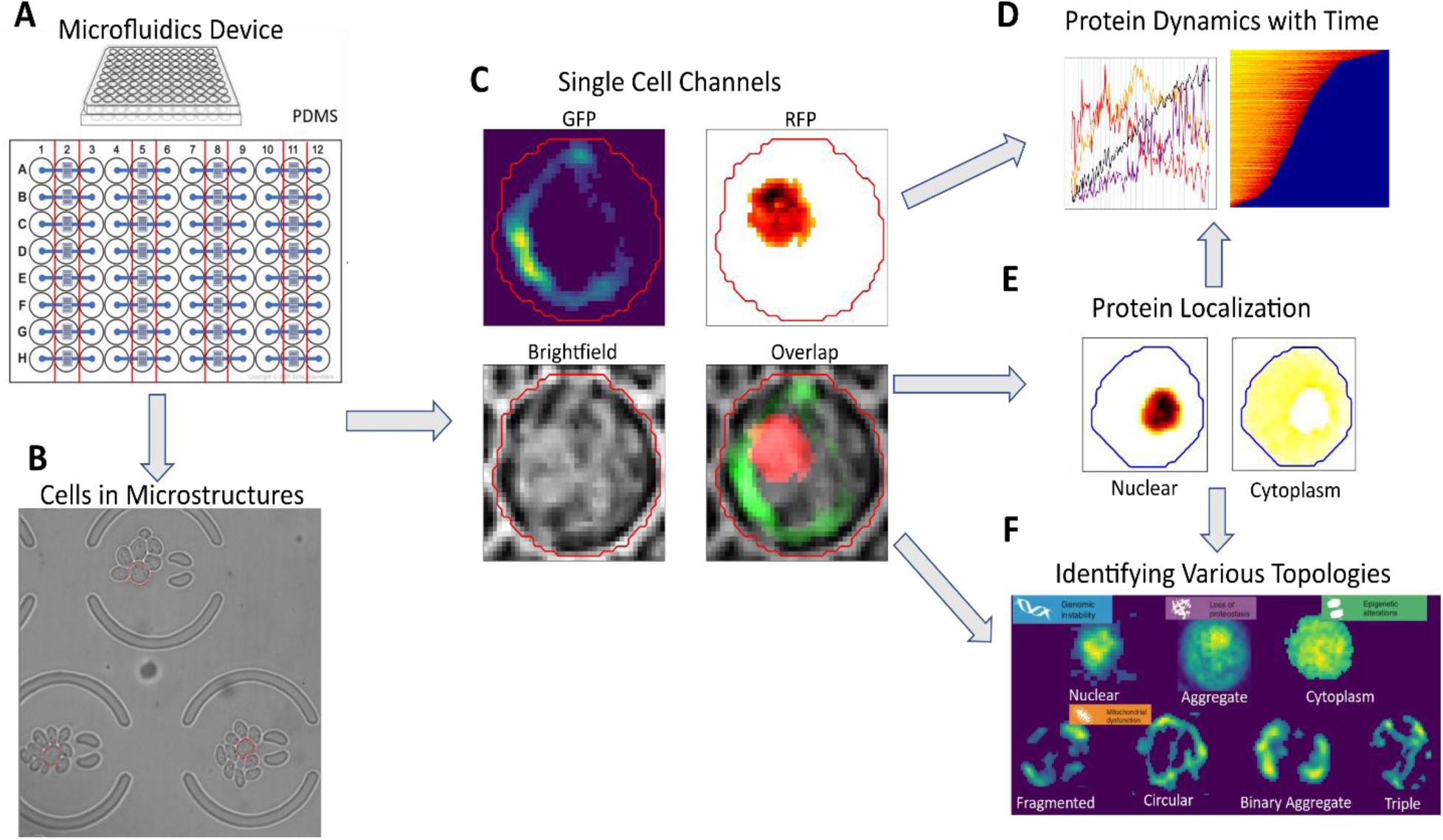
Tracking the dynamic trajectories of single cells: A high-throughput platform for single cell imaging and the pipeline for image analysis. (A) A microfluidic device with 32 independent modules, interfacing with a 96 well plate. Each module is aligned to three wells (e.g., A1, A2, A3), with the two side wells (e.g., A1 and A3) connected to the microfluidic device (depicted in blue) for liquid handling and cell loading. The center well (indicated by the red lines, e.g., A2) is aligned to the observational area of the device with ∼220 microstructures (see B); each can trap a single cell. (B) A brightfield image of yeast cells captured by a 63x oil lens. The red circle indicates the mother cells, and the large circular structures with two open ends are the microstructures used to trap cells. (C) Segmentation of a single mother cell with the corresponding fluorescent channels, GFP, RFP, Brightfield, and the overlap. (D) Quantification of protein dynamics over time using a python image analysis pipeline. (E) An example of how protein colocalization can be separated in segmented cells. (F) Topologies captured and classified by an AI model, using a mitochondrial marker (Cox4) as an example.

Using the DAP and the 96-well microfluidic device, we systematically analyze the dynamic trajectories of 34 strains each with a single molecular marker fluorescently tagged by GFP. These markers cover a wide range of biological processes, including the majority of aging hallmarks, including genomic instability, loss of proteostasis, mitochondrial dysfunction, epigenetic alteration, de-regulated nutrient sensing and several stress responses (see Table 1 for the list of the molecular reporters and the biological pathway they represent). To further assess the dynamic relations between these markers, we analyzed ∼34 strains with a pair of molecular markers tagged by GFP and mCherry respectively.

We performed time-lapsed imaging of single mother cells throughout their lifespan to track how the molecular marker change its expression and spatial distribution and extract other parameters (such as cell size, cell cycle time etc.) of the cells during the course of aging. Single cells are plated on a microfluidic device, after which cells are imaged using three channels brightfield, green, and red, with .4um Z stacks (Fig. 1B,C). Next using a tailored machine learning and AI model, time series data were generated to measure protein and phenotypic changes in individual dividing *S. cerevisiae* mother cells (from 20 to over 100 cells for each strain), over a span of 72 hours, with once every 20 min. Protein signal intensities, localization, and other phenotypes are then quantified in single cells (Fig. 1D-F). Subcellular localization of protein intensity is determined by a Gaussian Mixture Model (GMM). We validated GMM for determining subcellular localization using dual reporter strains with additional marker with known cellular localization (see below an example for nuclear localization). These single cell dynamic data allow us to calculate a number of parameters such as cell size, organelle size (such as the size of the nucleus), length of cell cycle, intensity and localization of the fluorescent signals, and topology of different organelles as a function of time.

### Identification of nuclear concentration of proteasome as a predictor for lifespan of individual cells

Since we have both the dynamic trajectory and the lifespan for each single cell, we can correlate some of the dynamic parameters with the lifespan, to identify potential lifespan markers. By simply correlating the expression level of a reporter at a given time with the lifespan of the cells that are still dividing at the time, we identified several potential lifespan markers, among them the reporters for proteostasis stands out (Fig. S1). Further analysis confirmed the proteosome concentration in the nucleus as a good predictor for the lifespan of individual cells (Fig. 2). Whereas, the cytoplasmic proteasome concentration weakly predicts lifespan of individual cells (Fig. S2).

**Fig 2.**
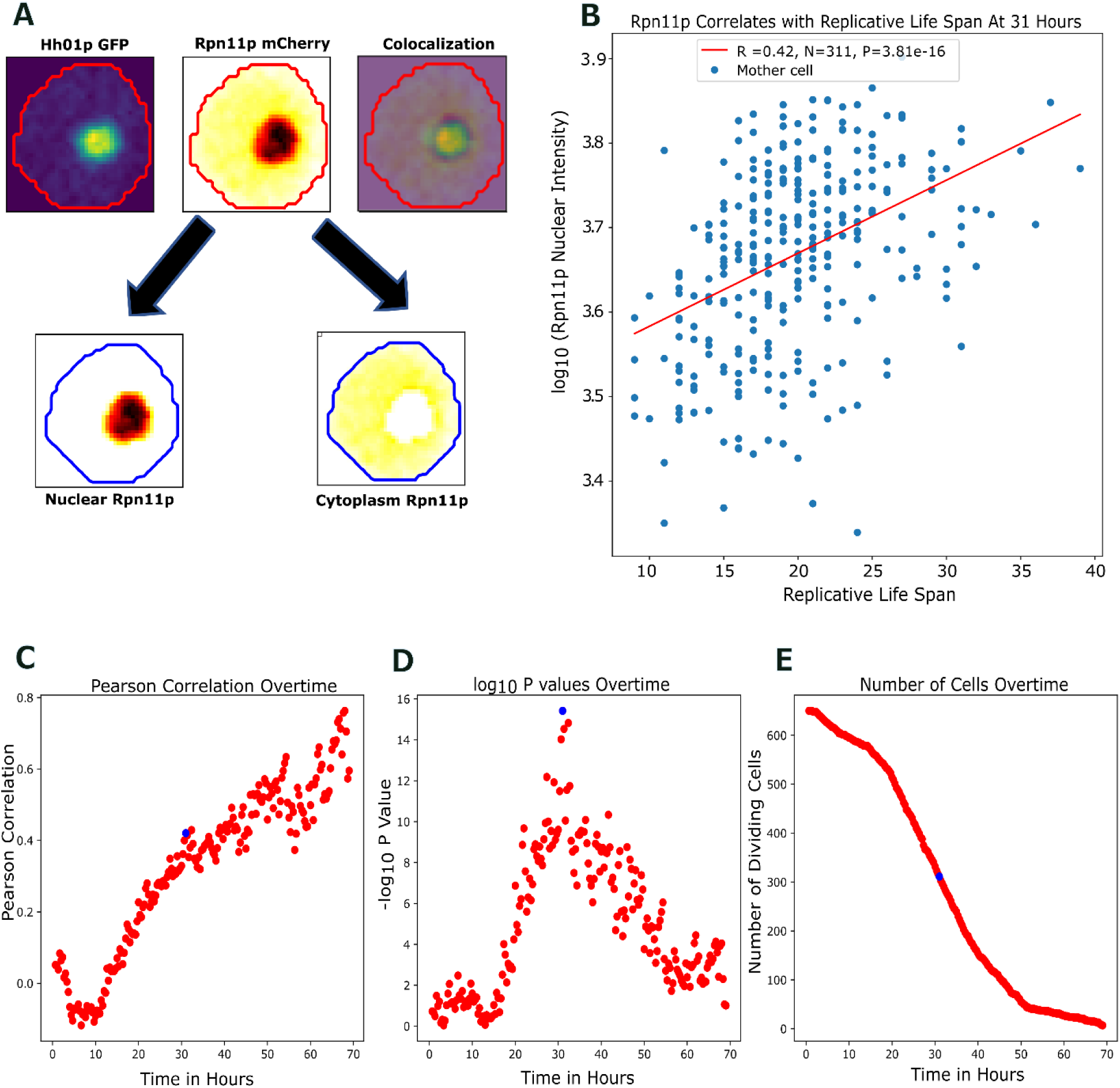
**Nuclear proteosome concentration at middle age correlates with the lifespan of individual cells**. (**A**) Representative images of a single cell with dual-reporters Rpn11p-mCherry and Hh01p-GFP. Colocalization is apparent in the image on the top right. A Gaussian Mixture Model (GMM) is employed to segregate and quantify regions based on Rpn11p intensity. This method facilitates the determination nuclear vs. cytoplasmic regions and protein localization. (**B**) Nuclear Rpn11p intensity at 31 hours strongly correlates with lifespan of individual cells. The x-axis displays the Replicative Life Span (RLS) of individual mother cells that are still dividing at the 31-hour mark. The y-axis exhibits the log10 intensity of nuclear Rpn11p at 31-hours. The Pearson coefficient is 0.42, with a total of 311 dividing cells. Statistical significance is given by a P value of 3.81e-16. (**C**) Pearson correlation coefficient between Rpn11p nuclear intensity and RLS as a function of time. Only cells that are still dividing at time t are counted. (**D**) the -log10 P value as a function of time for the observed correlation between Rpn11 nuclear intensity and RLS. (**E**) Number of cells that are still dividing at time t. The blue dots highlight the instances at 31 hours where the Pearson correlation P value reaches the maximum. The presented data aggregates findings from 12 independent experiments, each differing in terms of the GFP-tagged protein, but consistently featuring an Rpn11p- mCherry tag (total number of cells N= 650).

For the proteasome analysis, we used a dual reporter strain with Rpn11p with mCherry and Hh01p with GFP. Rpn11p is a subunit of the proteasome regulatory particle lid subcomplex (19S proteasome regulatory particle) and the proteasome has been documented to localize in both the cytoplasm and the nucleus of a yeast cell. Using GMM, we were able to differentiate between the nuclear and cytoplasmic regions of Rpn11p signal (Fig. 2A). The partitioning by the GMM is validated by the independent reporter Hho1p, which is a linker histone localized in the nucleus. There is a strong correlation between the segmented nuclear area using Rpn11p nuclear region identified by the GMM and the nuclear region indicated by Hho1p (Fig. S3).

We identified a strong correlation between nuclear Rpn11p intensity and replicative lifespan (RLS) for individual cells. The correlation R(t), calculated between Rpn11 nuclear concentration at a given time t and the lifespan across individual cells that are still dividing at time t, increase with time and reached maximum significance around 30 hours (Fig. 2B-D), when about half of the cells are still dividing (Fig. 2E). At later time points, as the number of dividing cells further decreases, the significance dropped. At the point of highest significance, a Pearson correlation coefficient of 0.42 is reached, giving rise to a P value of 3.81e-16. Cells with higher level of Rpn11p nuclear intensity tend to have longer lifespan (Fig. 2B), suggesting that cells with higher nuclear proteasome concentration are better at maintaining proteostasis. We observe only a weak correlation between cytoplasmic proteasome concentration and lifespan, but with a different sign (higher level correlates with shorter lifespan) (Fig. S2), suggesting that the nuclear proteasome plays a more important role in maintaining proteostasis in relation to lifespan.

### Nuclear proteasome concentration exhibits distinct dynamics from its cytoplasmic counterpart – relation to size scaling and nuclear size dynamics

The ability of nuclear proteosome concentration but not the cytoplasmic counterpart to predict lifespan prompted us to further analyze the dynamics of the proteasome in the two cellular locations in relation to the lifespan of individual cells. Examination of the dynamics of proteasome in single cells collectively revealed a striking difference: while the cytoplasmic concentration maintains more or less a steady level throughout much of the lifespan, the nuclear concentration reached a peak level around a characteristic time of ∼10 hours, and then rapidly decreases (Fig. 3 A, B). This feature is more apparent if we divide cells into long-lived and short-lived groups. Short-lived cells are classified as cells with a RLS less than 15 and long-lived is classified as cells with RLS greater than or equal to 15. While both groups reached their peak value around 10 hours, shorter lived group showed a more rapid decrease of the nuclear concentration than the longer lived group (Fig. 3 C, D).

**Fig 3.**
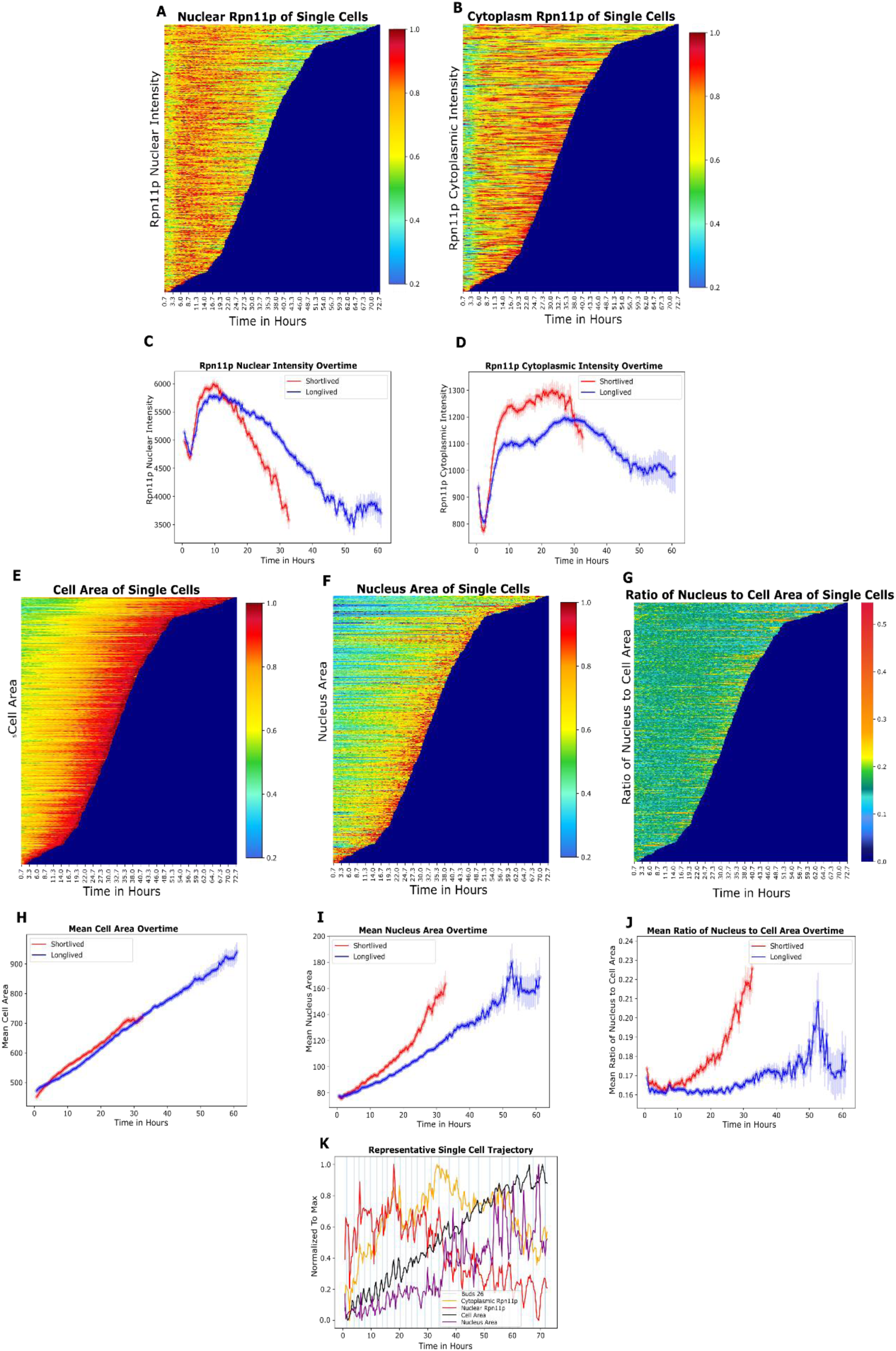
**Proteasome concentration in nucleus decreases more rapidly than that in cytoplasm during aging and correlates with the distinct dynamics of nuclear size vs. cell size**. (**A**) heatmap of Rpn11p intensity as a function of time in single cells. Each horizontal line represents a single cell. Cells are order by their lifespan from top (longest) to bottom (shortest). Rpn11 nuclear intensity is indicate by the color. (**B**) heatmap displaying the cytoplasm Rpn11p intensity for single cells as a function of time. In both A and B, for each single cell the intensity is normalized to the maximum value across the entire lifespan thus the range is between 0 and 1. (**C**, **D**) Rpn11 nuclear (C) and cytoplasmic (D) intensity averaged across single cells. Cells are divided into short lived and long lived groups, and the average is calculated separately for the two groups (red curves: short lived group; blue curves: long lived group). Short-lived cells are classified as cells with a RLS less than 15 (N=327, mean RLS=9.16) and long-lived is classified as cells greater than or equal to 15 RLS (N=323, mean RLS= 20.21). Data beyond the time points when the actively dividing cells become less than 25 are not shown (too noisy). (**E**) heatmap of the cell area as a function of time for single cells. (**F**) heatmap of the nucleus area as a function of time for single cells. For each single cell, cell area and nucleus area are normalized to the corresponding maximum across the lifespan. (**G**) heatmap of the ratio between the nucleus area and cell area as a function of time for individual cells. The ratio is calculated using the un-normalized sizes. (**H**) mean cell area of the short-lived (red curve) and the long-lived cells (blue curve). (**I**) mean nucleus area of the short-lived (red curve) and the long-lived (blue curve) cells. (**J**) mean nucleus area to cell area ratio for the short-lived (red curve) and the long-lived (blue curve) groups. Standard Error Mean (SEM) is the shaded region. Cells are taken from the aggregation of twelve independent experiments with a Rpn11p-mCherry tag (N= 650). (**K**) A representative single cell trajectory showing the dynamics of cell size (black), nuclear size (purple), cytoplasmic Rpn11 intensity (yellow), and nuclear Rpn11 intensity (red). The intensities are normalized to the maximum values. The cell is imaged every 20 minutes. The blue vertical line indicate the appearance of a bud.

Interestingly, the dynamic behaviors of proteasome seem to correlate with cell size and nuclear size dynamics. Both cell area and nucleus area individually correlated to RLS, reaching a Pearson correlation of -.42 and -.21 at time 27 and 11 hours respectively (Fig. S4A-F). The best correlation between nuclear size and lifespan (r = -0.42, P ∼10^(-18)) at 27 hours) is as strong as that between nuclear proteasome and lifespan (r = 0.42, P ∼ 10^(-16) at 31 hours). As the cells grew older, their cell size (as measured by the area) increased linearly with time, with the same slope for both the short-lived and the long-lived groups (Fig. 3 E, H).

We found that the nuclear size also increases with age, but in contrast to cell size, the longer-lived cells are better at maintaining a linear growth, while the short-lived cells display accelerated increase with a curvature (Fig. 3F, I). As a result, long lived cells maintain a constant ratio of nuclear size vs. cell size up till ∼ 30 hours while short lived cells display significance increase of the ratio after 10 hours (Fig. 3G, J). Interestingly, the ratio of the two organelle’s also weakly correlates with RLS (Fig. S4G). It has been observed in previous studies that the nucleus of different sized yeast cells maintain isometric size scaling, i.e., it scales proportionally to the cell size^19,20^. Our data indicates that during aging, the isometric size scaling is maintained early in life but broken later in life, and that long-lived cells are better at maintaining the scaling than short-lived cells. The rapid increase of nuclear size and the ratio of nuclear size vs. cell size seems to correlate with the rapid decrease of nuclear proteasome concentration, a relationship we further explore with a mechanistic model that utilizes single cell dynamic trajectories containing information of proteasome concentrations, cell and nuclear size, and cell division time. A typical example of the dynamic trajectories is shown in Fig. 3K.

To make sure that Rpn11 reporter faithfully represents proteosome expression level, we also tested another proteasome reporter Rpn6, which is an essential, non-ATPase regulatory subunit of the 26S proteasome lid. Rpn6 reporter recapitulated the same dynamic behavior observed with Rpn11 reporter (Fig. S5).

### A simple mathematical model of transport predicts decreased nuclear proteasome concentration due to nucleus size increase

To explore the relationship between proteasome concentration and nucleus size, we developed a mathematic model for the dynamics of nuclear concentration based on simple transport (Fig. 4A). Assuming that proteasomes are synthesized and assembled in cytoplasm and transported into nucleus, the difference of nuclear concentration ΔIn between time (t-1) and time t, is determined by the amount of transport across the nuclear membrane (the first term on the right hand side of the equation), degradation γ, and volume dilution due to size increase (second term on the right hand side of the equation). The transport term is proportional to the product of the rate of transport per unit area C1 and cytoplasmic concentration, with a scale factor of 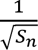 due to the different scaling behavior of volume vs. surface area.

**Fig 4.**
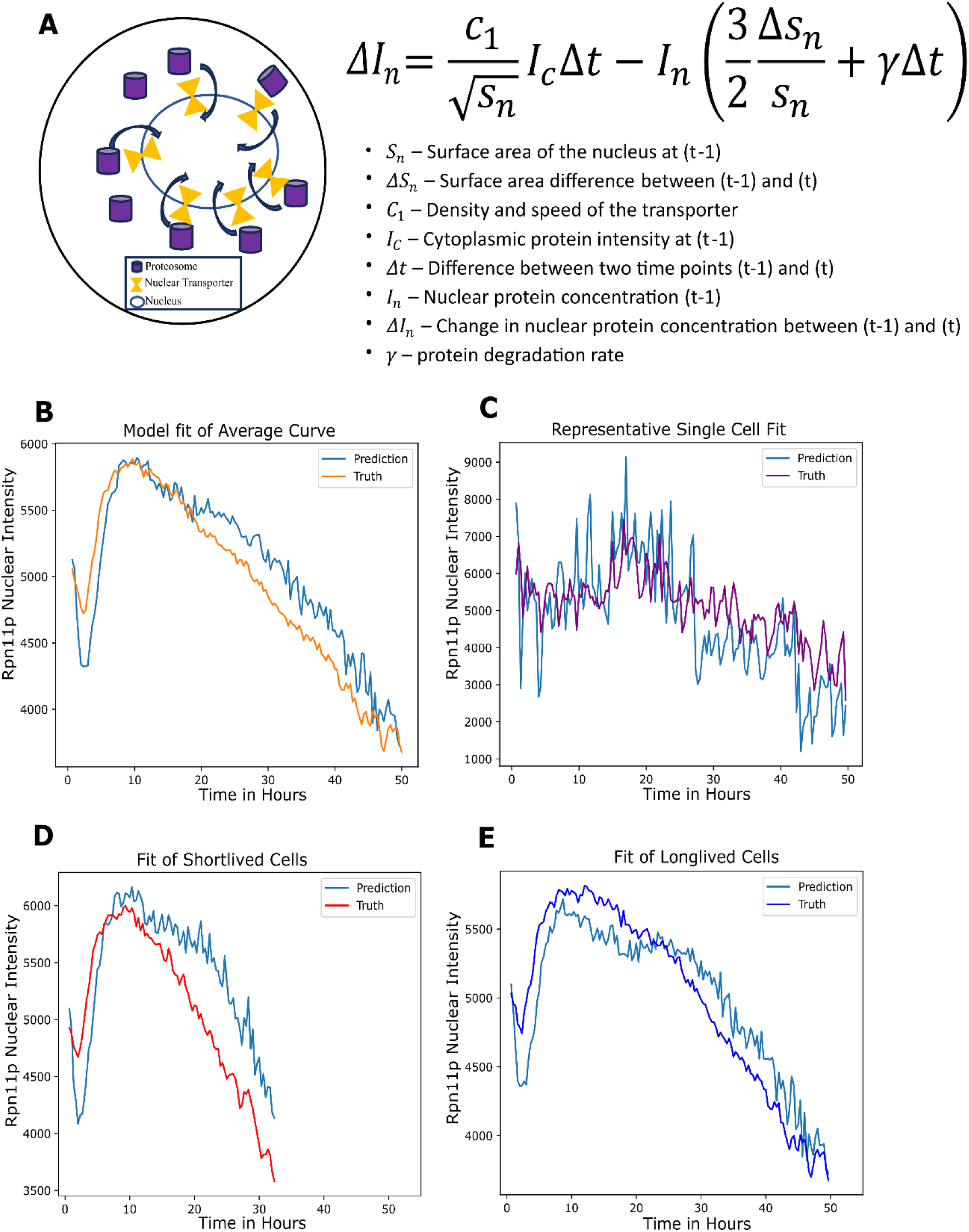
**A simple model of nuclear transport predicts nuclear proteosome dynamics from nuclear size dynamics**. (**A**) A schematic of the simple model of nuclear transport of proteosome (left) and the corresponding difference equation describing the dynamics (right). The model assumes: 1) proteosomes are synthesized and assembled in cytoplasm and transported to nucleus by transporters in nuclear membrane; 2) the transport flux per unit area is proportional to the product of cytoplasmic intensity and the transport efficiency per unit area (C1) which depend on both the density and speed of the transporters. For a derivation of the equation, see supplement notes. The meanings of the variables in the equation are explained below the equation. (**B)** The model described above predicts nuclear Rpn11 intensity using cytoplasmic intensity and nuclear size dynamics as inputs. Orange curve is the actual data (averaged nuclear Rpn11 intensity) and blue curve is the model prediction. (**C**) Model prediction of Rpn11 nuclear intensity as a function of time in a representative single cell. Purple curve is the actual data and blue curve is the model prediction. (**D**) & (**E**) Model prediction of Rpn11 nuclear intensity in short-lived cells group (D) and long-live cells group (**E**) using the same set of parameters (C1 and γ). Data beyond the time points when the actively dividing cells become less than 25 are excluded.

The volume dilution term is proportional to 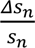 with a 3/2 factor similarly due to difference between relative surface area change vs. relative volume change (Supplementary Table 2). The unit of time in the model is set to be the time lapse between two consecutive imaging.

Using this simple model, we can predict the dynamics of nuclear concentration from that of the cytoplasmic concentration and nuclear size, with two tunable parameters C1 and γ. We found that the model can predict reasonably well the dynamics of nuclear concentration averaged over the individual cells (Fig. 4B), and with similar parameters also the dynamics of the individual cells (an example is shown in Fig. 4C, Supplementary Table 3). Moreover, the model also predicts the different dynamics for both the long-lived and the short-lived groups, with the same parameters. This model results suggest that the different dynamics between the long- and short-lived groups could be accounted for by their different nuclear size dynamics. Short-lived cells have accelerated nuclear size increase that breaks size scaling early, which can lead to more rapid decrease of nuclear proteasome concentration according to the model.

It is worth noting that according to the equation in Fig. 4A, nuclear concentration will decrease even if the cytoplasmic concentration is maintained as a constant, due to the 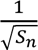 scaling factor in the input term. This is because the transport across nuclear membrane scale as R^2^ but the nuclear volume scale as R^3^, where R is the linear size of the nucleus; thus transport cannot balance degradation and dilution as the nucleus grows bigger and bigger. As a consequence the dilution of nuclear proteins can be a general phenomenon (see discussion).

### Time dependent over-expression of nuclear proteasome increases nuclear size and possibly decrease “transport efficiency”

To further test the model and to examine whether perturbing proteasome expression leads to change of cellular phenotypes (such as nuclear size) and lifespan, we engineered an inducible system to over- express Rpn4, a master transcriptional regulator that controls the transcription of all the proteasome components, at any specific time. The transcription factor Rpn4p is a short-lived protein (t1/2≤2 min) and is rapidly degraded by the proteasome and the ubiquitin ligase Ubr2p, giving rise to a negative feedback that makes the traditional over-expression difficult to achieve^21,22^. We utilized a truncated version of Rpn4 (Rpn4*), which has the degradation sequence removed (with the deletions Δ1–10 and Δ211–229) to cut off the negative feedback^22^. We placed Rpn4* under an estradiol inducible promoter system^23,24^. The truncated Rpn4 (Rpn4*) increases the proteasome activity, as measured by fluorogenic substrate, but does not affect cellular growth (Fig. S6 and Fig. S7). We picked two different time points for Rpn4* induction, identified through a non-linear Granger Causality method that indicated times at which the Rpn11p intensity is predictive of the nuclear area (Fig. S8). The timepoints are 5.5 hours, corresponding to a time early in the life of the cells, and 19.2 hours, corresponding to the mid-life where significant correlation between nuclear proteasome and lifespan has developed. Induction of Rpn4* at both points leads to significant increase of Rpn11p nuclear intensity (Fig. 5 A-B) as well as cytoplasmic intensity (Fig. S9A- B). Using the cytoplasmic proteasome concentration and nuclear size as input, the mathematical model was able to predict the induction profile for the nuclear proteasome concentration (as reported by Rpn11) for perturbations starting at both time points, by adjusting only a single parameter C1. We found that a smaller C1 is needed in order to fit the induction (C1 ∼25.8 ) compared to that of the control (C1 ∼30.8 ) (Supplementary Table 3 and Fig. 5 A, B). Together with the increased nuclear area and decreased transport efficiency, the model fits the induction of nuclear Rpn11p intensity which has a much smaller fold change (e.g, ∼0.3 fold increase for induction at 19.2 hours) compared to the fold change of cytoplasmic intensity (∼1.5 fold increase for induction at 19.2 hours) (Fig. 5B and Fig. S9B).

**Fig 5.**
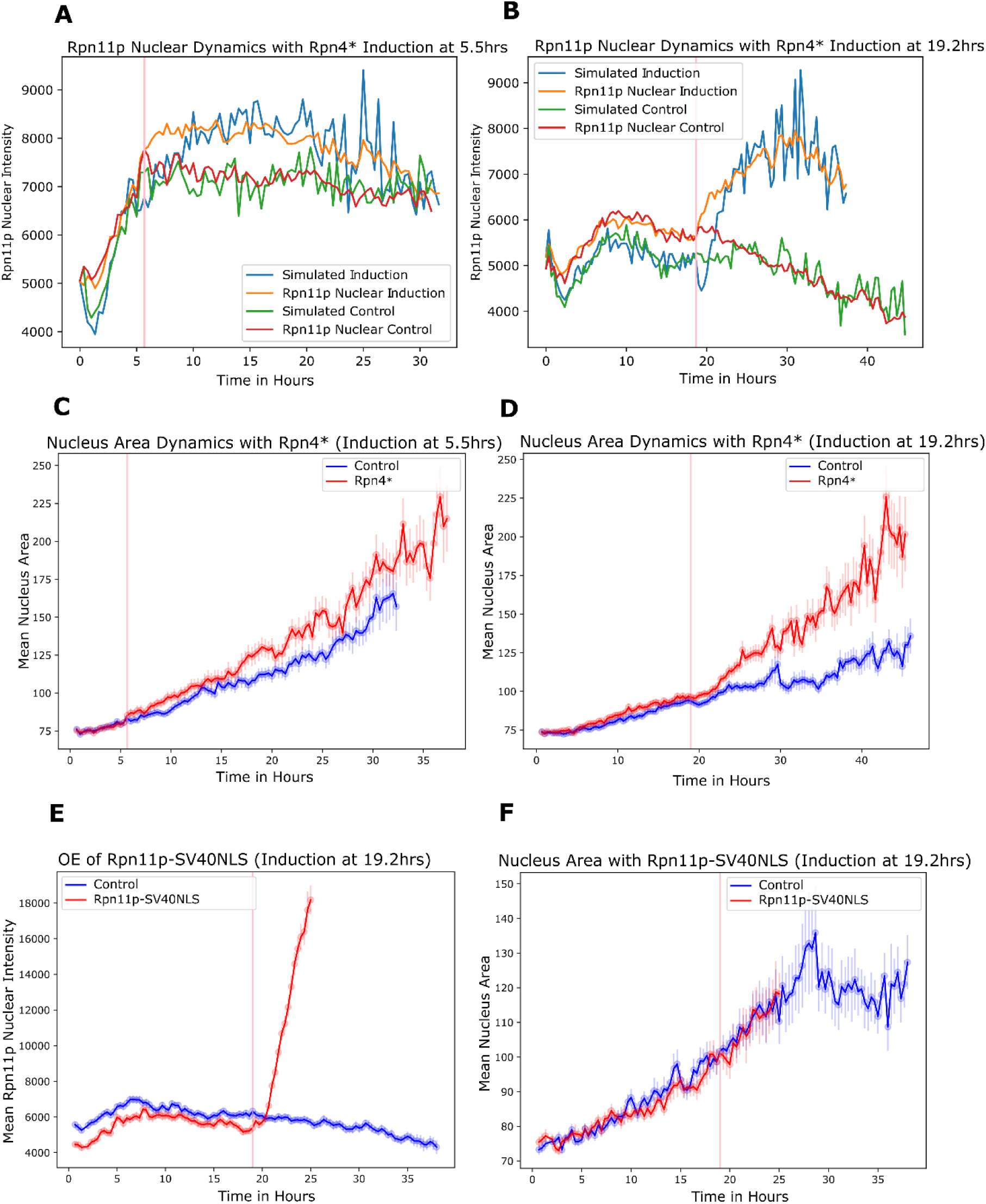
Time dependent over-expression of nuclear proteasome increases nuclear size. (**A & B**) Rpn11p nuclear intensity as a function of time from two different time dependent perturbation experiments. A truncated version of the transcription factor Rpn4 (a master regulator of proteosome) with the degradation sequence removed (called Rpn4*), is placed under an estradiol inducible promoter. Rpn4* was induced at 5.5 hours (**A**) and 19.2 hours (**B**) as indicated by the pink vertical lines. Rpn11p nuclear intensity was measured (orange) and fitted with the theoretical model (blue). Rpn11p nuclear intensity without the induction (control, red) and the corresponding model fit (green) are also shown. (**C & D**) the mean nucleus area as a function of time from the two time dependent perturbation experiments described in A and B. Red curves are for cells with Rpn4* induction and blue curves are controls without induction. (**E & F**) Rpn11p nuclear intensity (**E**) and nucleus area (**F**) as a function of time with over- expression of Rpn11p-SV40NLS induced at 19.2 hours (indicated by the pink vertical lines). Red curves are for cells with the induction and blue curves are controls.

Interestingly, Rpn4* induction led to an increase in nucleus area thereby supporting the prediction based on the non-linear Granger Causality approach (Fig 5. C-D). This change of the nuclear size is not an artifact due to the increased influx of nuclear Rpn11p, as over-expressing Rpn11 itself using a Rpn11p- SV40NLS tag (with a nuclear localization signal) increased Rpn11 nuclear intensity by more than 3-fold but did not change nuclear area (Fig. 5E-F). We did note that endogenous levels of Rpn11p were sequestered to the cytoplasm when Rpn11p-SV40NLS tag was induced (Fig. S9C).

Together these experiments and model analysis suggest that over expressing proteasome leads to increased nuclear size and decreased rate of transport per unit area as inferred from the model; both can serve as negative feedback to maintain nuclear concentration of the proteasome.

We did not observe significant difference in terms of lifespan between Rpn4* induction and the control. Possible reasons are rescuing nuclear proteasome is not sufficient, or the dosage of the induction is not optimal (see discussion).

### Inferring the dynamic relationships between aging hallmarks: proteasome vs. mitochondria

Aging is characterized by a set of molecular hallmarks^25,26^; the majority of these hallmarks are exhibited during yeast replicative aging^2^. While the individual hallmarks have been extensively studied, the temporal order of and inter-relationships between these hallmarks are not well understood. Part of the motivation for us to study the dynamic trajectories of single cells with pairs of reporters is to see if we can infer the temporal order and perhaps even the causal relations between different hallmarks. Since the dynamic behavior of proteasome is tied to lifespan of individual cells, and previous studies indicated that loss of the proteasome activity affects mitochondrial activity, we decided to explore the dynamic relationship between proteasome and mitochondria ^16,27,28^.

Since mitochondria displays complex morphological features, we developed AI based phenotype classifier to characterize the dynamics of mitochondria (see methods). This phenotype classifier allows us to classify the topology of mitochondria as well as the number of foci present in single cells (Fig. S10).

The topologies are classified at a coarse-grained level into three categories: “circular”, “aggregate”, and “other”. The “other” category includes many morphological structures that cannot be classified as either circular or aggregate (Fig. 6A and Fig. S10B ).

**Fig 6.**
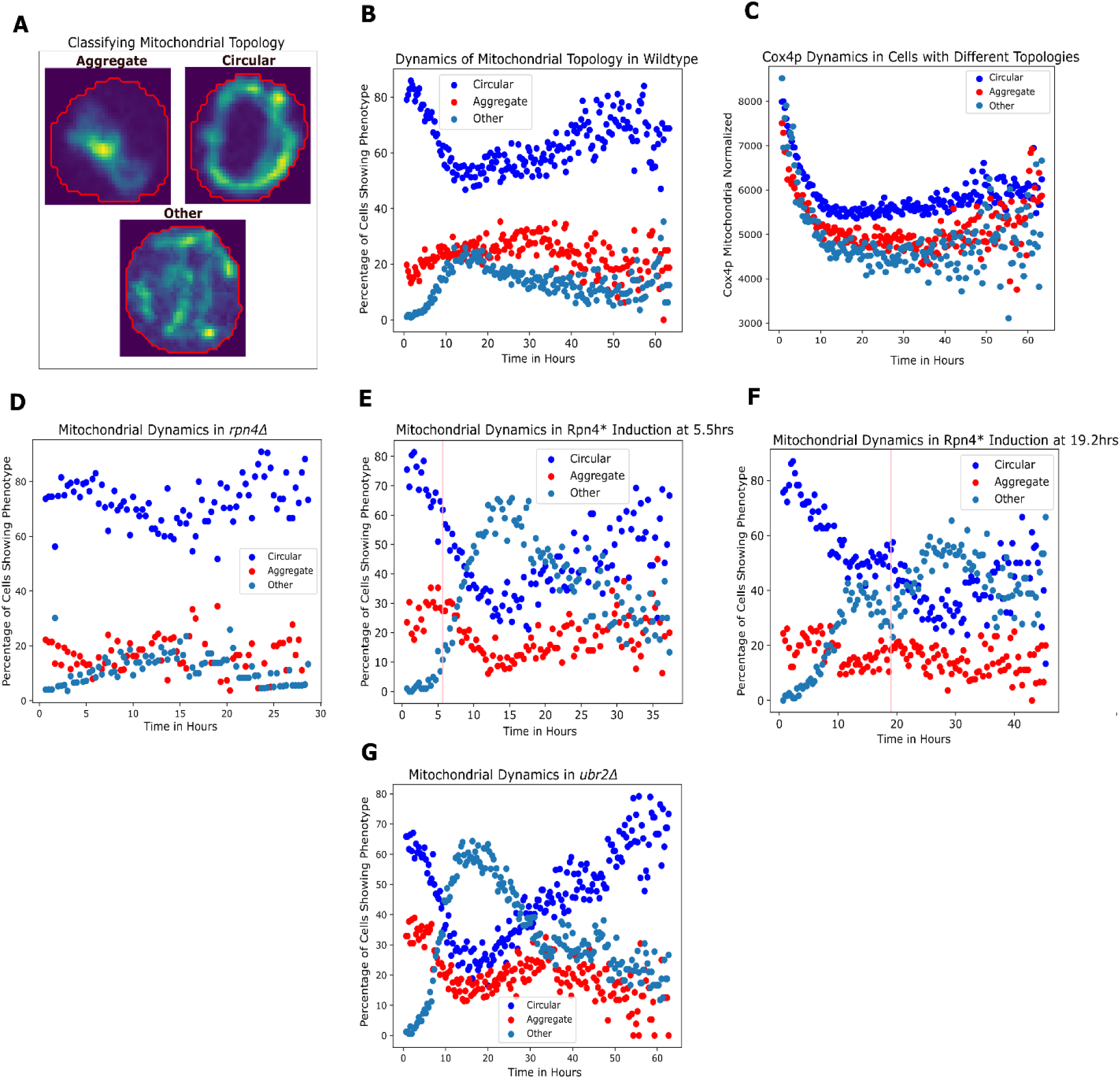
Dynamic perturbation of proteosome leads to change of mitochondrial morphology. (**A**) Example images of mitochondria morphologies classified as aggregate, circular, and other. (**B**) Percent of cells with different mitochondria morphologies (circular: blue; aggregate: red; other: light blue) as a function of time in the wild type strain. (**C**) Average mitochondrial Cox4p intensity (normalized by the cell area) for subgroups of cells with different mitochondrial morphologies. (**D**) Percent of cells with different mitochondria morphologies as a function of time in the *rpn4*Δ mutant strain (proteosome decreased). (**E & F**) Percent of cells with different mitochondria morphologies as a function of time with Rpn4* induction at 5.5hours (E) and at 19.2 hours (F). Pink vertical lines indicate the time of induction. (**G**) Percent of cells with different mitochondria morphologies as a function of time in the *ubr2*Δ mutant strain (proteosome increased). Data for all the subfigures are aggregate of two or more experiments performed independently.

The probability for a single cell to be in each of these states changes over time. Thus, we get a dynamic trajectory of the mitochondrial state for each single cells. The percentage of cells displaying circular morphology first decreases and then increases as the cells get older (Fig. 6B), while the other two categories showed opposite trend. Interestingly, we found that circular morphology may have better respiratory function as indicated by increased Cox4p activity relative to the other two topology categories (Fig. 6C). Cox4p is the subunit IV of cytochrome c oxidase which is the terminal member of the mitochondrial inner membrane electron transport chain. This is consistent with previous report that as yeast cells get older, their metabolism shifts from fermentation to respiration^29,30^. When cells are separated based on whether they spend the majority of time in the circular or aggregate state, we found that cells displaying the circular morphology lived longer than cells displaying the mitochondrial aggregate morphology (Fig. S11A-B). Additionally, the budding rate between the morphology was also different with cells of circular morphology having slower budding rate than those with aggregate morphology (Fig. S11C-D).

To determine whether proteasome perturbation affects mitochondrial morphology, we followed the dynamics of mitochondria using Cox4p as the reporter while perturbing the proteasome activity. Several studies used Cox4p to measure mitochondrial activity such as cellular respiration^31,32^. We found that deletion of *rpn4Δ*, which lead to a decrease in the proteasome, resulted in an increase in the “circular” mitochondrial state (Fig. 6D). Mitochondrial Cox4p intensity was also higher in *Δrpn4* (Fig. S12A-C) suggesting that decreased proteasome leads to increased mitochondrial respiration.

We next temporarily induced the proteasome by over-expressing Rpn4* at times 5.5hours and 19.2 hours Interestingly, temporal overexpression of the proteasome led to an increase in the “other” state and decreased in the “circular” state (Fig. 6E,F). Similarly, the overexpression of the proteosome in the *ubr2Δ* (the ubiquitin ligase that helps degrade Rpn4) deletion mutant increased the “other” state and decreased the “circular” state (Fig. 6G). Temporal overexpression of the proteasome led to a minor decrease in Cox4p intensity, similar to *ubr2Δ* deletion (Fig. S12D-L). Thus, perturbing the proteosome could influence mitochondrial morphology and activity, and our data suggests that increased proteosome leads to decreased mitochondrial respiration.

We next set out to determine if changing mitochondrial morphology could affect the proteosome by analyzing the *dnm1*Δ strain. Dnm1 is a dynamin related GTPase and has a role in mitochondrial fission^33^. Knocking out Dnm1 affects mitochondrial morphology and leads to aggregate mitochondrial structures (Fig. 7A, compared to the wild type behavior Fig. 6B)^33^. We confirmed that *dnm1*Δ affected both mitochondrial morphology and Cox4p intensity (Fig. 7 B-C). However, the dynamics of Rpn11p nuclear and cytoplasmic concentration remained largely unchanged (Fig. 7 D-E). Additionally, we did not see a major change in the nucleus area trajectory (Fig. 7F).

**Fig 7.**
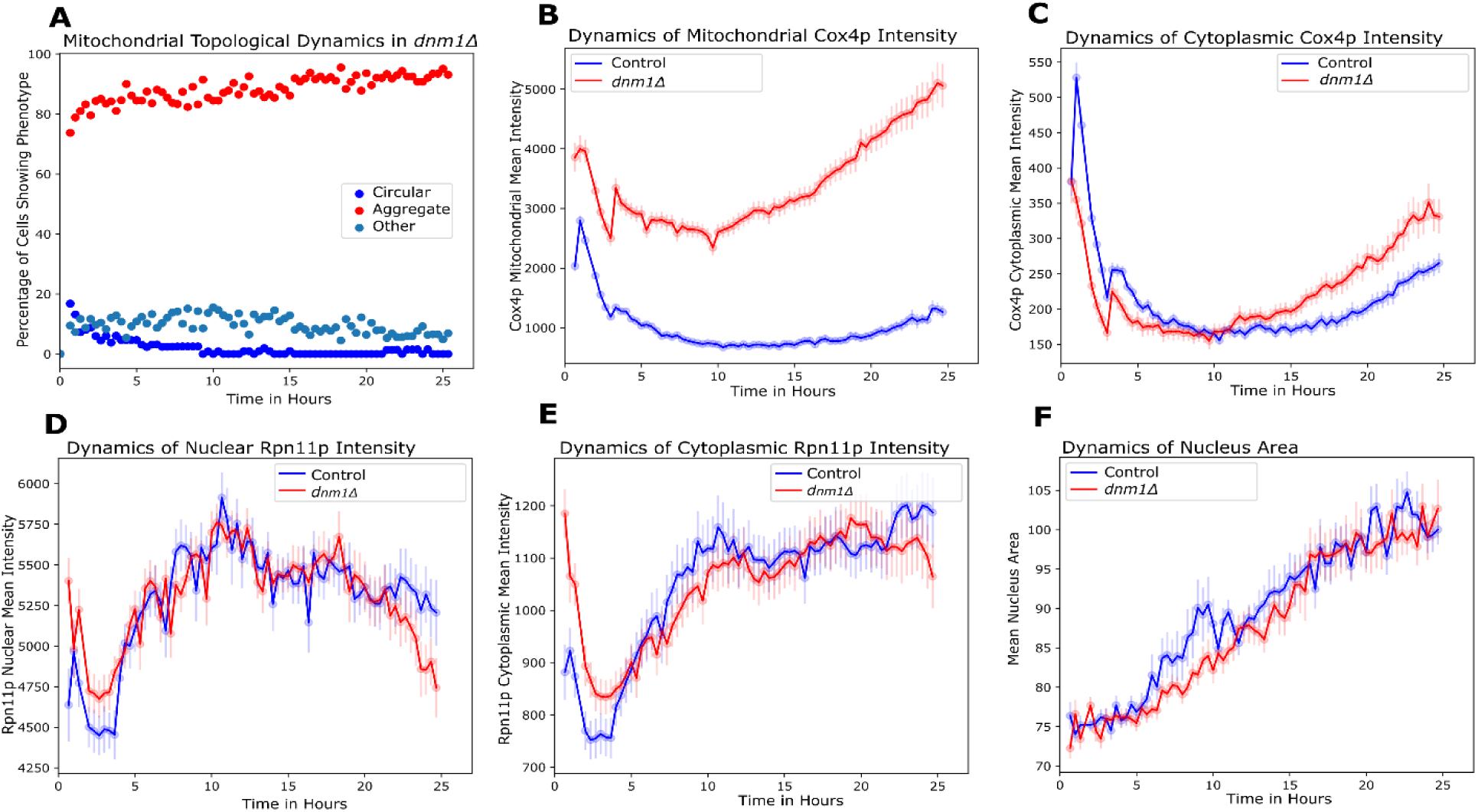
Perturbing mitochondrial morphology does not change proteasome dynamics or nuclear size. (**A**) Percent of cells with different mitochondria morphologies (circular: blue; aggregate: red; other: light blue) as a function of time in the *dnm1*Δ mutant strain. (**B -F**) Comparison between the wild type (blue) and the *dnm1*Δ mutant (red) strains of average mitochondrial Cox4p intensity (normalized by the cell area) (**B**), average cytoplasmic Cox4p intensity (**C**), average Rpn11p nuclear intensity (**D**), average Rpn11p cytoplasmic intensity (**E**), mean nucleus area (**F**).

Together, these data suggest that change of proteasome during aging may be upstream of change of mitochondria activity and morphology, potentially placing the loss of proteostasis upstream in the causal chain of events that drive the other aging hallmarks.

## Discussion

Yeast replicative aging is cell autonomous and can be studied in a constant environment in microfluidic device; thus it is an ideal model to analyze the process of aging from a dynamic systems perspective.

Using an engineered strain of yeast with inducible program to stop daughter cells from budding (the daughter arresting program, or DAP) and a 96-well plate formatted high-throughout microfluidic device, we monitored the dynamics of thousands of single mother cells throughout their lifespan with a broad range of molecular reporters. Our study indicates that loss of proteostasis maybe a major driver for aging. Out of the 34 reporters we analyzed, several reporters covering proteostasis (Hsp104p, Rpn11p, Trx2p, Vma1p, and Uba1p see Fig. S1) are predictive of the lifespan of individual cells. Analysis of dynamics revealed an interesting interplay between the dynamics of cell size, nuclear size, and nuclear proteasome concentration. As the mother cells age, their cell size (surface area) increase linearly with time. Initially nuclear size increase in proportion to maintain isometric size scaling. As the cells become older, their nuclear size increases faster than linear and isometric size scaling breaks down. At the same time, nuclear proteosome concentration decreases, and nuclear proteasome concentration at midlife is predictive of the lifespan of individual cells. These observations suggest that loss of proteostasis in the nucleus may be an important limiting factor for the lifespan.

We observed distinct dynamic behaviors for cytoplasmic proteasome concentration and nuclear proteosome concentration; the later declined much more rapidly with age. We showed that such dynamic behavior can be accounted for by a simple mathematical model of transport, taking into account the increase of nuclear size over time. This model predicts that even if the cytoplasmic concentration is kept constant, the nuclear concentration will still decline as the nuclear size increases. This is because transport scales proportionally to the surface area of nucleus (assuming the density and efficiency of the transporters do not change), while dilution and degradation scales proportionally with the volume, leading to an effectively input term that decreases with the increase of nuclear size (Fig. 4A and Supplementary note). This is purely a geometric property of scaling of surface vs. volume, and thus as a mechanism for the dilution of nuclear factors should be general. We expect that other nuclear factors that need to be synthesized in cytoplasm and transported into nucleus will be similarly diluted as nuclear size increases, unless an additional feedback regulation is invoked. We did observe that another nuclear factor Uba1p, which is an Ubiquitin activating enzyme (E1), exhibited similar behavior that can be explained by the same mathematical model (Fig. S13). Thus, potentially dilution of multiple nuclear factors can limit lifespan. This would explain why over-expressing proteasome (by over-expressing RPN4*) is not sufficient to extend lifespan. It also suggests that interventions that lead to better control of nuclear size or feedback control of nuclear pore complex (e.g, increasing the density) may be a better strategy for lifespan extension. This model may be relevant to aging in other higher eukaryotes, as increase of nuclear size was also observed in mice and human cells (Fig. S14)

From a dynamic systems perspective, yeast aging reflects a dynamic system that drifts gradually to become unstable, leading to the eventual catastrophic failure. One obvious drifting parameter is the cell size. We observe that cell surface area increases linearly with time, and that the rate of increase is the same for different cells. The reason for this constant increase is unclear. We hypothesize that for every bud produced, there is a fixed amount of newly synthesized materials pumped into mother cell membrane/cell wall – the excess amount over what is needed for a new bud. It has been observed in different contexts that cell nucleus scales isometrically with cell size^19,34^. Our analysis indicated that even if different organelles scales appropriately with cell size, it is still difficult to maintain homeostasis if the cell size increases constantly. E.g., for proteins needed both in cytoplasm and in nucleus (such as proteasome), if their concentration in cytoplasm needs to be kept at a constant, their nuclear concentration will decrease due to the increase of nuclear size; thus overall their total amount will scale sub-linearly with the cell size. Indeed, in recent studies of size scaling of mRNA and proteins, a set of proteins are found to scale sub-linearly with cell size, and they were enriched for nuclear proteins^35^ . Interestingly, proteome changes with size seems to correlate with proteome changes due to aging, suggesting that an important component of age induced changes may be caused by cell/organelle size change^35^.

In addition, increase in cell size can lead to an effective decrease of nutrient influx from the environment, by the same geometric argument applied to nuclear proteasome, assuming there is no particular feedback regulation of different nutrient transporters. The hypothesis that increase of cell size can lead to decrease of effective nutrient influx is consistent with the observation that with aging yeast seem to elicit stress response and undergo diauxic shift where metabolism shifted towards respiration^29,30^, which we also observed with the classification of mitochondrial morphology (Figure 6B). We thus hypothesize that an early driver of aging could lead to the increase of cell size and drive the loss of homeostasis. Interestingly cells that rigorously control its size (such as E. coli and fission yeast pombi) do not seem to show obvious aging phenotypes^36,37^.

Cellular aging is a complex phenomenon where multiple processes become dysregulated. Our study suggested that by simultaneously tracking the dynamics of multiple molecular reporters in naturally aging cells, it is possible to infer the temporal order and even the causal relationships between different dynamic processes. We provided an example where change of proteasome activity proceeds the change of mitochondria topology (and possibly function), and experimentally tested the causal relation by perturbing one process and following the response of another. In particular, we showed that over- expressing proteasome leads to decreased circular morphology and decreased mitochondrial respiration.

This general approach can be extended to other molecular reporters to infer the temporal order and potentially causal relations between them, and hopefully resolve whether multiple processes degenerate in parallel during aging or there is a cascade of events that drive aging. This is an important issue as for the latter scenario, early interventions at the upstream of cascade can be effective at slowing down or even reversing aging.

Our study produced large scale single cell dynamic data that can serve as a useful resource for the aging research community to analyze the dynamics of other markers and potentially infer causal relations between them. It is also a useful resource for training and testing AI based models that identify early dynamics features that are predictive of lifespan, and may be good targets for longevity interventions. In addition, the observed dynamics trajectories of a large number of single cells can be used to build and test dynamic systems based models with the basic principle applicable to more complex multiple cellular organisms.

## Author contributions

MM and HL conceived and designed the study. MM performed and led all experiments and bioinformatic analysis. CD created the microfluidic device as well as the original DAP strain and provided guidance for experimental design. MM performed AI training and image analysis. JZ created a user-friendly interface to visualize single cell trajectories as well as provided bioinformatic and image analysis guidance. MM and HL analyzed data. HL developed the mathematical model. MM and HL wrote the paper.

## Methods

### Strains

A modified By4741 (*MATa his3*Δ*1 leu2*Δ*0 met15*Δ*0 ura3*Δ*0*) was used to create the Daughter Arresting Program (DAP). The DAP yeast strain was generated by integrating an exogenous nucleic acid sequence upstream of the start codon of the PMA1 gene, encoding an essential plasma membrane protein. This construct consists of two opposing promoters: (1) a mother-specific promoter driving an exogenous copy of PMA1, and (2) a conditional promoter regulating the endogenous PMA1 gene. The exogenous PMA1 can be tagged with a Red Fluorescent Protein (RFP) or infrared fluorescent protein (iRFP). This can assist in helping researchers track the mother cells. For the purposes of this study iRFP channel was not used to identify the mother cells. The PMA1 integration site was verified via PCR and Sanger sequencing. To induce arrest, the DAP strain is switched from galactose rich media to glucose media, thereby inhibiting the daughter cells.

### Cell Tracking and Segmentation Pipeline

For segmentation a pretrained U-Net model was trained to receive one channel 512x512 images. The pretrained model was trained on a curated dataset containing only segmented yeast mother cells. The purpose was to teach the model to differentiate between cells and the microfluidic device wells. The pretrained model was then retrained on a training set data composed of the YeaZ dataset and our hand segmented dataset of yeast cells. The YeaZ dataset consists of high-quality segmented yeast microscopy images, encompassing over 10,000 cells. It includes diverse conditions such as wild-type cells, mutants, stressed cells, and time-lapse sequences^38^. Cell masks are generated by turning all values, including Yeaz dataset into 0 or 1. Binary Cross Entropy with logits loss function and Adam Optimizer is used with a starting learning rate of .001. The model was trained for 100 epochs on batches of 5 with 755 images, each image contained 3-300 cells. Out of the 755 images, 423 are images made from experiments using the Li lab’s microfluidics device, while the rest are composed of the YeaZ dataset. The model’s accuracy was determined on a testing dataset of 14 images, which were created through a separate imaging experiment that the model has never seen, each image contained more than 5-100 cells. Augmentations used included random horizontal flipping, random vertical flipping, random rotations (0,24,90,135,270), gaussian blur, random gaussian noise, colorjiter, CutMix, Random Crop, and translations. The final Unet model is trained to receive a single 512x 512 image. Out of the 17 Zstacks taken per time for each well and experiment, only the top two brightfield images were extracted and merged. The two sharpest brightfield images for each mother cell are identified using a Laplacian absolute value mean, which would select for the sharpest Z stack per mother cell. Merging more than two of the brightfield images leads to bad segmentation and blurred images. Images are then normalized to values of 0-1 and fed to the trained Unet model to determine cells. After segmentation, the boundary of the cell was determined by using skimage find_contours function. Yeast mother cells were identified during the first timepoint of imaging and tracked using a python-based pipeline. Should issues arise due to segmentation, the pipeline used a watershed-based approach to further help separate out daughter cells from mother cells. The cell’s previous cell eccentricity and cell area is used to determine whether segmentation proceeded correctly. If not, the model would back track to create a better segmentation. If the model failed to track the cell, manual hand curation is utilized. This occurred in cases where large shifts in the microscope occurred or several mother cells are plated relatively close, pushing the others. To filter out any outliers, a Pearson correlation is computed between the cell area vs. time (as cell area should increase linearly with time).

Only cells with a Pearson value greater than .7 are kept for downstream data analysis. The segmentation pipeline is robust in being able to track cells regardless of whether they are DAP.

### Segmentation of Nucleus in Liver Cells

The liver data are shared through Diana Jurk at Mayo Clinic. The nucleus segmentation procedure was executed through the Python programming language utilizing multiple packages: readlif.reader, scipy.stats, time, numpy, skimage.measure, and pandas. The process was initialized by loading images from LIF files using the LifFile function from readlif.reader. The nucleus segmentation process was initiated by obtaining the DAPI-stained image of the cell nucleus. A binary mask of the nucleus was generated through adaptive thresholding, where pixels with intensities greater than the mean plus one standard deviation were set to true. Post thresholding, any remaining holes in the binary mask were filled using the binary_fill_holes function from scipy’s ndimage module, and small objects less than 700 pixels in size were eliminated using the remove_small_objects function from the skimage.measure module.

Using a similar method the cytoplasmic mask was created. For each nucleus, we made sure that it was kept within the cell boundary by using a cytoplasmic marker. Only nucleus sizes that shared one overlap with a cytoplasm label are kept. Further filtering included removing cells where the nucleus area is larger than the cytoplasmic area. Cells are then checked by eye to determine proper nucleus segmentation.

### Segmentation of Cryo-Section Lung Data

The mice lung data are shared through Soledad Reyes De Barboza Nabora at UCSD and Tiang Peng at UCSF. The U-Net architecture was utilized for segmentation of cellular structures in microscopic images, beginning with a pretrained U-Net model designed for 512x512 yeast segmentation. This model was trained on a subset of nucleus data, augmented with additional images from various sources. The dataset included 81 images from a study published on the European Bioinformatics Institute website (https://www.ebi.ac.uk/biostudies/bioimages/studies/S-BSST265), 34 images from previously published aged fibroblast DAPI stains, and 18 miscellaneous images of lung fibroblast and passaged fibroblast cells. Two hand-annotated images were used for validation, these images had more than 100 cells each. All images had reasonably segmented cells. The U-Net model was trained with a dataset that was augmented through random rotations, zooms, shifts, CutMix, and flips. Brightness and contrast of the images were randomly adjusted for additional variety. Given the usage of a pretrained U-Net model, only 30 epochs were required for training with a starting learning rate of 0.001. The Adam Optimizer and a Binary Cross Entropy with Logits Loss were used in the training process. A smaller batch size of 3 was favored for better learning. Images larger than 512x512 were patched and fed to the model for training. For segmentation, if the image exceeded the size of 512x512, it was cropped to patches of this size. Padding is added if the image is too small. A threshold selecting for nucleus areas between 80 and 500 was applied. This filter eliminated most double nuclei and overlapping cells. To validate the findings of the violin plot, more stringent thresholds were applied using eccentricity of .6 and .7 to filter out cells that were less circular. The general trend of older cells possessing a larger nucleus area was maintained. The figure focused on using the nucleus area threshold of 80 to 500 as it captured most of the cells.

### Foci Segmentation

For the main training set data, 628 files, focusing heavily on mitochondrial foci, were utilized. A total of 179 images were used to facilitate model training, supplemented by 13 images of Cox4p-GFP and 16 of Rpn11p-mCherry foci that were augmented by hand. A separate hand-curated validation set contained 25 files from an independent Cox4p-mCherry-tagged experiment not included in the training set data.

Augmentation techniques such as CutMix, random erasing, horizontal and vertical flipping, random rotations, translations, gaussian blur (sigma of 2), and random gaussian noise were implemented to enhance model robustness. The weights were initialized using the yeast U-Net model, designed for 512x512 input images and segmentation. However, the image size for training and foci detection was reduced to 40x40. Cells were identified using our segmentation pipeline, and the mother cells were cropped. To maintain a consistent 40x40 image size, padding was added and blank areas were filled with the minimum intensity value found within the cropped cell. For model training, we employed the Binary Cross-Entropy with Logits Loss (BCEWithLogitsLoss) function, Adam optimizer, and an initial learning rate of 0.0001. The model underwent 50 epochs of training with a batch size of 25.

### Phenotype Classifier

Initially we started with seven phenotypes, classified for a training set data of 10713 images and validation dataset of 965 images. Phenotypes are identified using GFP tagged data collected from our single reporter strains. Phenotypes are easier to see when performing a background subtraction on each of the 17 zstacks before aggregating the zstacks together. Training images and validation images are identified through a Mean Maximum Discrepancy Variational Autoencoder to identify novel protein characteristics and confirmed by hand. The phenotypes included Circular (2022 images), Aggregate (2338 images), Cytoplasm (1549 images), Other (1475 images), Fragmented (1357 images), Binary Aggregates (1144 images), and Triple (828 images). The Other phenotype included images of phenotypes not identified. The phenotype labeled Aggregate can be used to differentiate between Hsp104p aggregates, nuclear localization, and mitochondrial aggregates. The Circular, Fragmented, Binary Aggregates, and Triple phenotypes pertain specifically to the mitochondria but could be used on other organelles. The model was tested on two separate datasets to determine its accuracy. The mitochondrial marker was Mdm38p-GFP and the cytoplasmic was Trx2p-GFP treated with rapamycin, for a total of 965 testing images with about 120-150 images per category. To create a classifier model, we used a pretrained ResNet50 model trained on the ImageNet database. The ResNet50 is then retrained on our phenotype data. Here, the first layer of the pretrained ResNet50 model were merged into one channel. All portions of the model were frozen except for the first conv1D and the last linear layer. Segmented single mother cells are fed to the model, these cells are resized to a 224x224, to utilize the pretrained properties of the ResNet50 model. Augmentations used: random rotations (0,45,90,135,180,270), horizontal flipping, vertical flipping, gaussian blur, random noise, unsharp filter, and random translations. To further help the model identify key differentiating phenotypes, a random threshold is performed along with downsizing and then resizing the images to increase blurriness. The MixUp augmentation did not help with model performance. Model was trained with a batchsize of 32 for 100 epochs with a Adam Optimizer and a starting itineration rate of .0001. Accuracy is further tested by using knockout experiments to determine if the model could accurately identify phenotypes. The *dnm1*Δ is known to cause mitochondria aggregation and ablate the mitochondrial circular phenotype. The trained ResNet50 model corroborated this.

For the statistical analysis presented in the figures, we grouped seven phenotypes into three to increase the number of cells in each category: Circular, Aggregate, and Other. The Fragmented phenotype is re- classified as Circular. The Binary and Triple phenotypes are re-classified as Aggregate, and the rest as Other.

### Microfluidic Setup

Cells were grown overnight in yeast extract peptone (YEP) medium supplemented with 10% galactose and 10% dextrose (YEP-GD: 2% peptone, 1% yeast extract, 10% galactose, 10% dextrose) to allow for balanced carbon metabolism and conditional promoter regulation. The following morning, cultures were diluted to an OD₆₀₀ of 0.2 (1:5 dilution) in fresh YEP-GD and incubated at 30°C with shaking (200 rpm) for 3 hours and 30 minutes to ensure continued growth. Cells were then washed three times in yeast extract peptone (YEP) medium (2% peptone, 1% yeast extract, no carbon source) by centrifugation (3000 × g, 3 min) and resuspended in YEP. Cells were incubated for 1 hour in YEP to induce temporary cell cycle arrest before microfluidic loading. After incubation, cells were diluted 1:2 in yeast extract peptone dextrose (YEPD) medium (2% peptone, 1% yeast extract, 2% dextrose) and loaded into a microfluidic chamber using a 20 PSI pressure-driven flow system for controlled deposition. 135 µL of fresh YEPD was added to the wells, followed by a continuous wash for 2 minutes at 0.5 PSI using a microfluidic pump to remove un-trapped cells and residual media. After washing, the microfluidic system was equilibrated by adding 280 µL of YEPD to one side of the well and 135 µL of YEPD to the other, ensuring uniform media distribution. To maintain chamber hydration, neighboring empty wells received 250 µL of sterile water.

### Confocal Imaging Protocol

Cells are imaged every 20 minutes using the lowest laser settings possible to view tagged proteins. Cells are incubated at 30 Celsius. A Zeiss Axio Observer Z1 at 63x oil objective is used to capture cells every 20 minutes and 17 zstack images are acquired with .45um distance. A Zeiss microscope was used with two laser settings 405nm and 588nm. Laser power was at 15% and exposure time was 200ms.

### Protein Quantification

Proteins were quantified using custom python code to identify single mother cells and quantify fluorescence. Median background level is obtained per zstack by observing areas without cell or well occupancy. Proteins are quantified by taking the pixel sum values per zstack of the region of interest designated by the mother cell segmentation program. Next the median background for that zstack is subtracted from the values. Protein intensity is determined by normalizing the total pixel values by the area of occupancy of the respective proteins or organelle areas.

### Gaussian Mixture Model (GMM)

To strengthen intensity differences among subcellular structures, the 17 z-stacks were first combined, which boosted signal to noise ratio. A Gaussian filter (σ = 3.5) was then applied to this aggregated image to help accentuate regions with distinct intensity while smoothing noise. A threshold was used to classify pixels at or above the cutoff (defining a high-intensity region), while those below were grouped as low intensity regions. From these GMM-defined regions, we quantified pixel intensity, total intensity, and total area. Visual inspection of these segmented regions allowed us to discern different organelles, including the nucleus, mitochondria, and vacuole.

### Induction Experiment Protocol

Fresh media containing 16nM of estradiol is made each time. Experiments were paused and old media was removed on both well sides. Fresh media of 135ul of new media were added. The well that would be induced, received fresh estradiol. The control would receive fresh YEPD media without estradiol. The media would be pumped using a PSI of .5 for 2 minutes. After which the respective media would be removed and replaced with a new fresh media of YEPD with 16nM of estradiol to induce overexpression or YEPD without estradiol. Volume added is 280ul for one side of the well and 135ul on the other side to ensure a proper pressure gradient. Experiments would resume ∼15-20 minutes after the initial addition of regular media and estradiol added media.

### Protein Tagging with Fluorescent Tags

*Saccharomyces cerevisiae* strains were generated using standard yeast transformation techniques to enable protein quantification and phenotypic analysis. Three fluorescent protein tags were employed for this purpose: Green Fluorescent Protein (GFP), mCherry, and a near-infrared fluorescent protein (iRFP). These tags were fused to the C-terminus of a target protein of interest, which was selected based on its well-characterized expression pattern and biological significance identified through our preliminary analysis. Selection markers for the fluorescent tags used are nourseothricin, Ura3, and hygromycin B. Plasma membrane P2-type H+-ATPase, Pma1p, is fused to iRFP to tag the individual mother cells, however, in our experiments the marker was not used to identify or keep track of the single mother cells.

### Gene Deletion

Kanamycin gene was inserted between the promoter and the terminator of a gene of interest. For homologous recombination, Primer designs of 57bp of the start of the coding sequence and 57bp of the end of the terminator are used. Deletion is confirmed through gel PCR and sanger sequencing composed of primers 500bp upstream and downstream of the gene of interest. For all knockouts at least 5 or more colonies are seen 3-4 days after plating cells on a kanamycin plate.

### Estradiol Plasmid Construction

The estradiol Z4EM plasmid was gifted by the Hana El-Samad lab^23,24^. The plasmid was remade to insert into the Ura3 region with a kanamycin selectable marker, site insert, and Cys4 protein. The Ret2 promoter is used to induce the expression of the activating protein. The Ura3 region was chosen as a site of genomic insert as it was previously indicated to have relatively small epigenetic changes with time. Cys4 protein was included in the plasmid in case multiple genes needed to be overexpressed. For this paper only one gene was expressed at a time during induction experiments. To insert a gene of interest, upstream primers containing a Kozak sequence for the start of the coding sequence along with the enzyme AVRII were constructed and the terminator sequence 200bp at the end of the gene was added with a enzyme sequence SBFI. If a gene had a fluorescent tag the terminator past the GFP tag was taken. To insert a gene, restriction digest enzymes AVRII and SBFI were used. Inserts were checked through gel PCR and sanger sequencing. Induction can be measured through live microscopy and is visually seen within 15- 20 minutes of estradiol induction. Flow cell cytometry also confirmed rapid induction and stabilization.

### Proteasome Activity Assay

Cells were cultured to OD600 ∼0.8 to 1.0 in 15ml of galactose rich media. Cell pellets were collected by spinning samples down at 1500RPM for 5 minutes and washing in 10ml of deionized water twice. Pellets were then stored in -80 for one night before resuspending in 50μl of cold lysis buffer [50 mM tris-HCl (pH 7.5), 0.5 mM EDTA, 5 mM MgCl2, and protease inhibitor cocktail tablet; Roche], and p200 was used to squirt cell mixture into a 2.0ml tube. The capped tube was then placed in liquid nitrogen, creating frozen droplets within the tube. Eight 3.0 mm Diameter beads (D1032-30) were then placed inside the tube with frozen droplets. At this point tubes are kept in liquid nitrogen. Cell breakage was performed by mixer milling. Mixer milling was performed three times at a frequency of 30 Hertz for 1 minute, samples and rack were placed back in liquid nitrogen for two minutes between breakage times. At this point the frozen droplets appear as a refined powder. 15μl of cold lysis buffer was then added to the tubes and samples were allowed to thaw on ice for 15 minutes before supernatant was collected at 13,000 rpm for 8 min at 4°C. Protein concentration was determined by a Bradford assay (Bio-Rad) using bovine serum albumin as standard and 3-5μl of samples. Proteasomal chymotrypsin-like activity was measured by using Suc-LLVY-AMC (Bachem), and proteasomal caspase-like activity was measured by using Ac-nLPnL- AMC (Bachem). Each reaction volume was 200μl, including 50μg of total lysate protein, 100μM peptide substrate, and lysis buffer. The fluorescent intensity was recorded after incubation for 15 min at 30°C in a TECAN with an excitation wavelength of 380nm and an emission wavelength of 460nm. Samples were recorded in the absence or presence of 50µg/ml of the proteasome inhibitor MG132.

### Temporal Ordering with Granger Causality

The primary intention is to determine whether feature 1 is predictive of feature 2 at a later time. The Granger Causality is expressed through a combination of Decision Trees and F-test in this analysis. Two datasets representing feature 1 and feature 2 activities at distinct time points were used. Both datasets were aligned based on a common time frame and cell population. The data points were normalized by Singular Value Decomposition (SVD) to equalize the scale. Moreover, the data for each cell were normalized to the 0-1 range to facilitate trend recognition. A Decision Tree Regressor was employed for the Granger Causality test. Granger Causality is generally used for linear relationships; however, the use of a Decision Tree Regressor allows us to capture the nonlinear relationships between datasets. The trees were optimized by setting the maximum depth as a function of the square root of the number of live cells in the sample, which adapts the model complexity to the data size. Furthermore, the Friedman Mean Squared Error criterion was applied as the function to measure the quality of a split. Time points with less than 56 live dividing cells are excluded. These predictions were then compared with the actual future feature 2 activities. The difference between predicted and actual feature 2 activities were calculated and subjected to an F-test, where the null hypothesis is that the variances of the two sets of differences are equal. The essence of the approach is to capture which model has less variation. The one that has less variation would thereby be a better predictor. The result of this F-test determines the Granger causality: if the p-value of the F-test is below a given significance level (alpha), it indicates past feature 1 activity has statistical significant effect on the future feature 2 activities, and hence we can infer that feature 1 predicts feature 2. Finally, the log10-transformed p-values (signed according to the sign of the F statistic) of the F-tests were assembled into a matrix where rows and columns represent different time points. A rolling window approach was applied with a window size of two where the maximum value within each window was selected. This helped to reduce the noise and increase the signal. Magnitudes less than two standard deviations of the signal greater than zero were turned to zero. Following the thresholding step, a gaussian filter with a σ = 1.2 was used to further smooth and enhance true feature distinctiveness. This matrix was then visualized using a heatmap, providing a visual overview of the temporal relationship between feature 1 and feature 2 across all time points. The predictive relationships were identified through negative logarithmic p-values, indicating statistically significant Granger causality from feature 1 to feature 2. A greater magnitude of these values represents stronger evidence of causality.

### Code Availability

https://github.com/takamichael?tab=repositories

