## Supplemental Figures for "Yeast aging from a dynamic systems perspective: Analysis of single cell trajectories reveals significant interplay between nuclear size scaling, proteasome dynamics, and mitochondrial morphology"

A

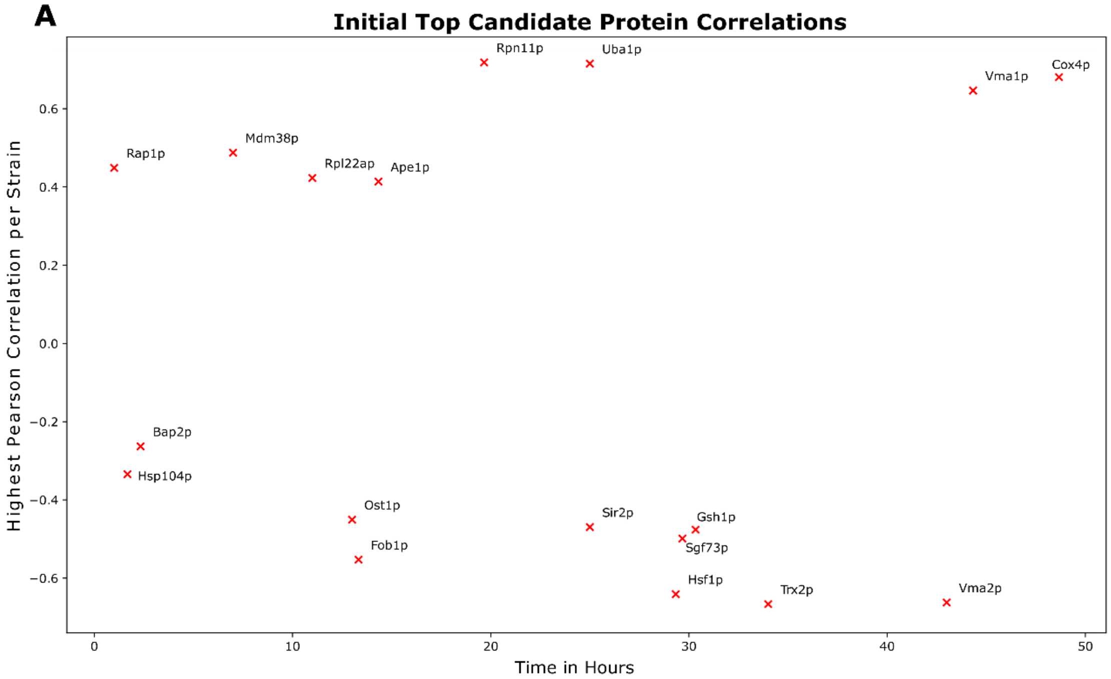


B

| Proteostasis Related Proteins | Pearson Correlation | Time in Hours |
|---|---|---|
| Hsp104p | -0.35 | 2 |
| Rpn11p | 0.7 | 22 |
| Uba1p | 0.68 | 25 |
| Hsf1p | -0.63 | 29 |
| Trx2p | -0.65 | 35 |

**Supplementary Fig. 1. Proteostasis related molecular markers are most predictive of lifespan**. (**A**) The scatter plot displays the top 19 protein vs. lifespan correlations identified from a mini-screen of 34 GFP-tagged markers. Proteins with extreme phenotypes, poor signal quality, or insufficient sample sizes were excluded. Each red "X" represents a distinct protein, tagged at its C-terminal with GFP, for which Pearson correlation analysis was performed. The x-axis denotes the time (in hours) at which the maximum Pearson correlation with replicative lifespan (RLS) was observed. The y-axis represents the Pearson correlation coefficient at the time. A minimum of 20 dividing cells was analyzed for each correlation measurement, though sample size varied by strain. (**B**) A table summarizing notable proteostasis-related proteins that exhibited significant correlations with RLS. The Pearson correlation coefficient and corresponding time point (hours) at which the maximum correlation was observed are reported for each protein.

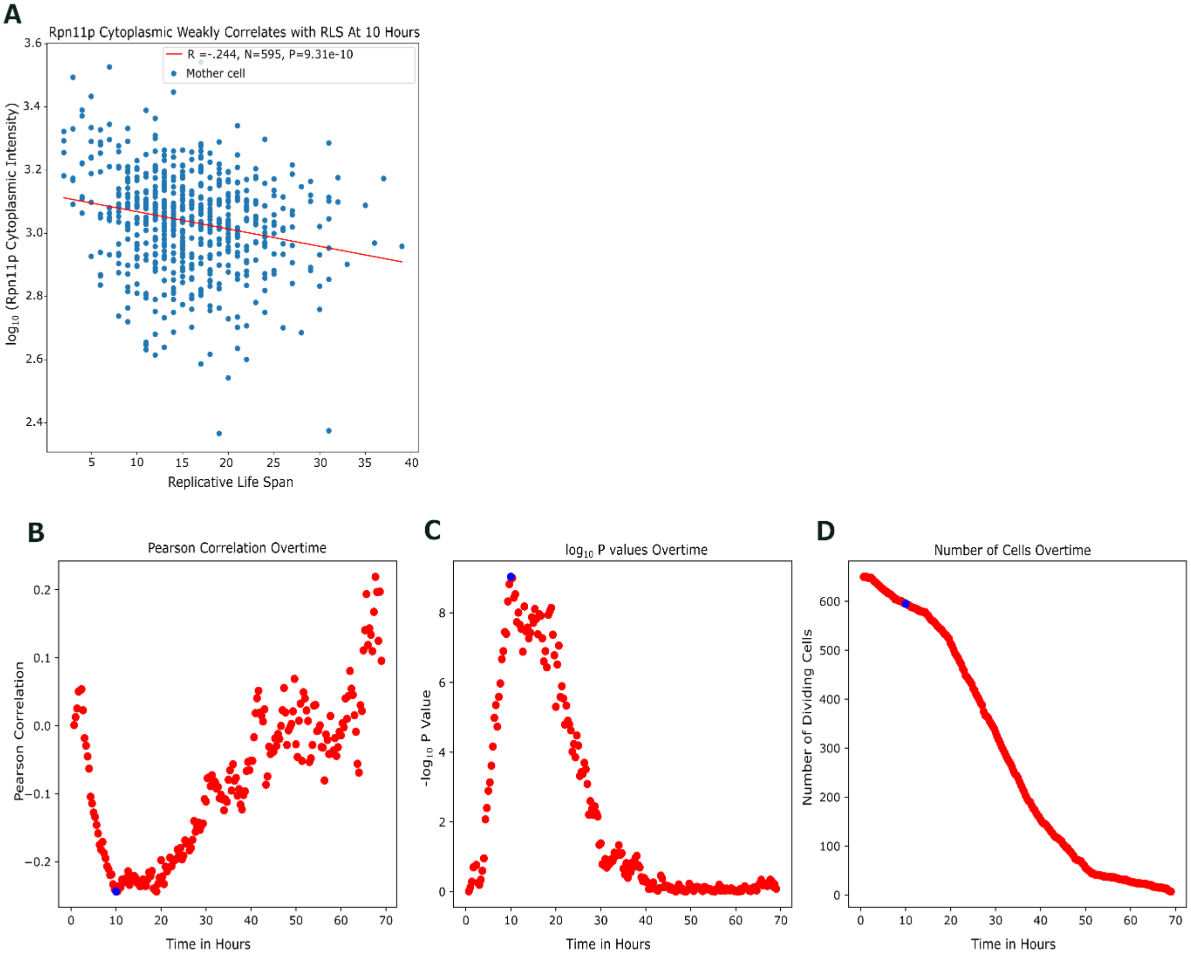


**Supplementary Fig. 2. Cytoplasmic proteasome concentration weakly correlates with lifespan**. (**A**) Scatter plot depicting the relationship between Rpn11p cytoplasmic intensity and replicative lifespan (RLS) at 10 hours. The x-axis represents the RLS of individual mother cells that were actively dividing at the 10-hour mark, while the y-axis shows the $\log_{10}$-transformed cytoplasmic intensity of Rpn11p at the same time point. A Pearson correlation coefficient of -0.244 was observed (N = 595 cells), with a p-value of 9.31e-10. The red regression line represents the trend in correlation. (**B**-**D**) Aggregated Rpn11p-mCherry data compiled from 12 independent experiments (N = 650 cells). Each experiment involved a different GFP-tagged protein, while Rpn11p-mCherry was consistently used as a reference marker. Only time points with ≥25 dividing cells were included in the analysis. (**B**) Pearson correlation of Rpn11p intensity over time. The x-axis denotes time (hours), and the y-axis represents the Pearson correlation coefficient between Rpn11p cytoplasmic intensity and RLS. (**C**) Statistical significance of correlation over time. The x-axis represents time (hours), while the y-axis shows the - $\log_{10}$-transformed p-values for the correlation at each time point. The highest statistical significance occurs at early time points, reaching the peak at ~10 hours, and gradually decreasing over time. (**D**) Number of dividing cells over time. The x-axis represents time (hours), while the y-axis shows the total number of live dividing cells at each time point. The blue dot marks the 10-hour time point corresponding to panel (A).

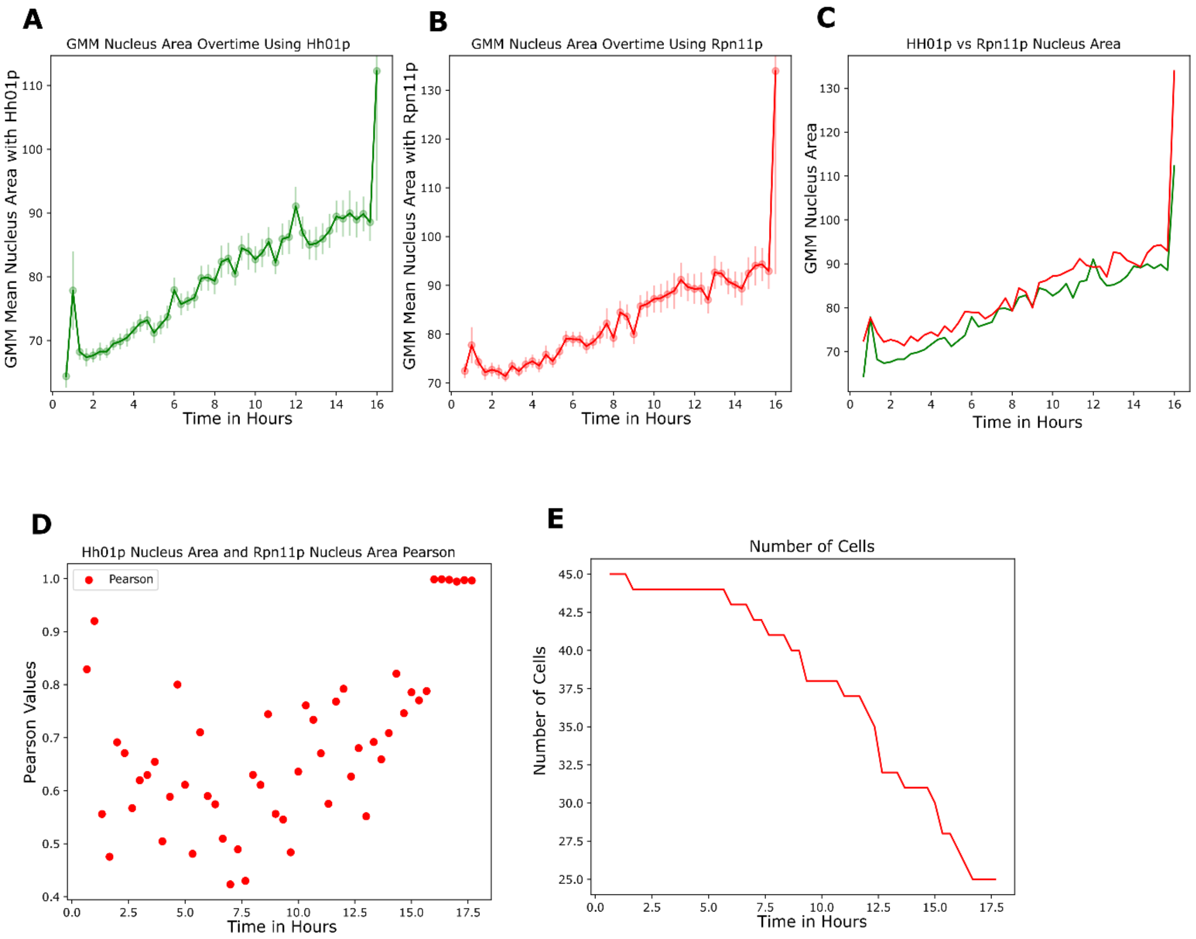


**Supplementary Fig. 3. Nuclear areas derived from Rpn11p reporter and Hho1p reporter are highly correlated**. (**A-B**) Mean nuclear area over time for Hho1p (green) and Rpn11p (red), respectively. The nuclear area was measured using a Gaussian Mixture Model (GMM)-based segmentation. The x-axis represents time (hours), while the y-axis represents the mean nuclear area (with standard error bars). (**C**) Comparison of nuclear area dynamics for Hho1p and Rpn11p. The nuclear area for both proteins is plotted over time, showing a similar trend in nuclear growth and segmentation. (**D**) Pearson correlation between Hho1p and Rpn11p nuclear area over time. The x-axis represents time, while the y-axis shows the Pearson correlation coefficient between the nuclear areas of the two proteins across single cells. (**E**) Number of dividing cells tracked over time. The x-axis represents time (hours), while the y-axis shows the number of actively dividing cells included in the analysis at each time point. The number of cells decreases as the experiment progresses. A total of 45 cells are tested.

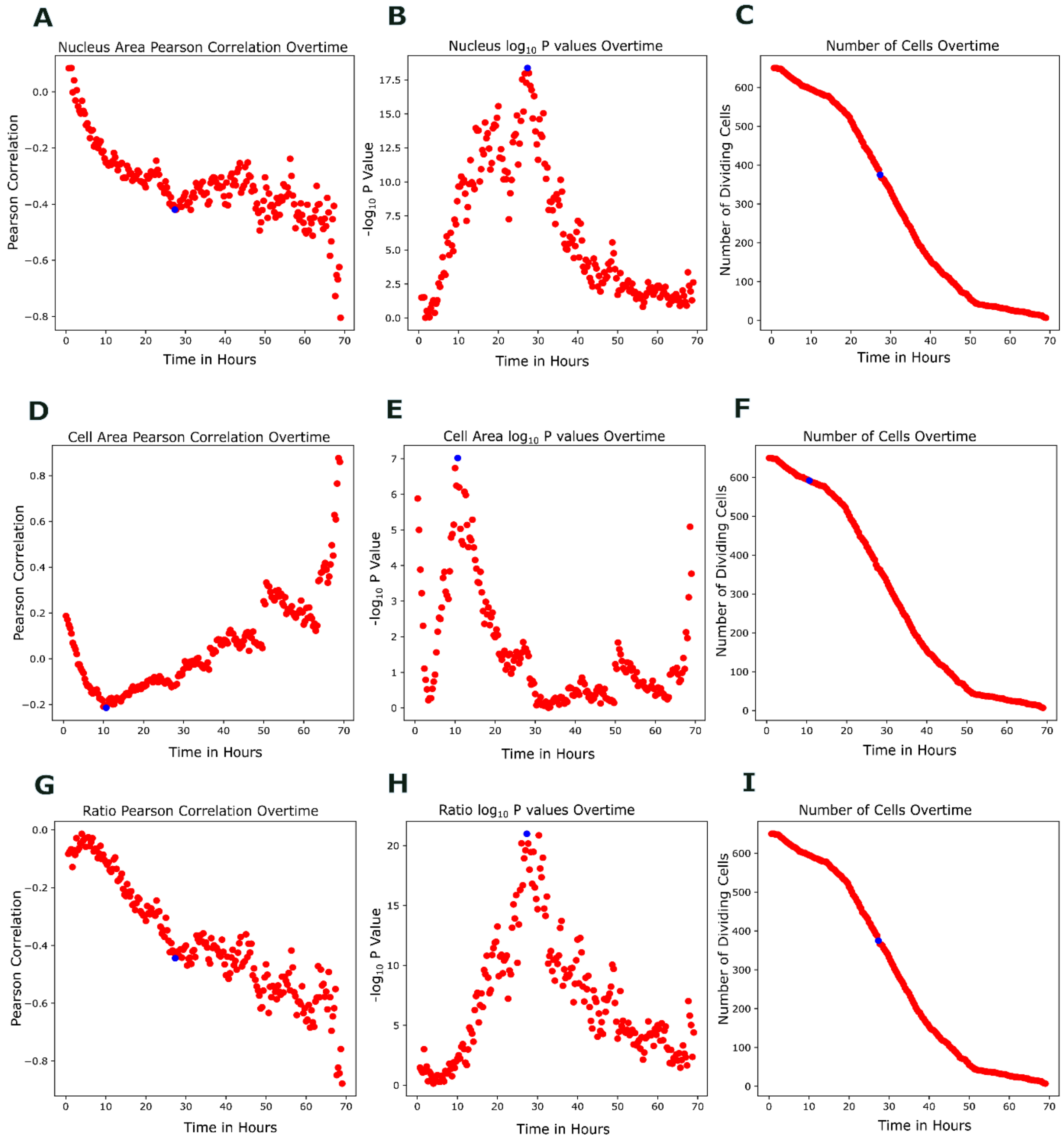


**Supplementary Fig. 4. Correlation between cell area or nuclear area vs. lifespan.** This figure presents Pearson correlations and statistical significance analyses of nuclear area, cell area, and their ratio vs. lifespan over time using nuclear Rpn11p-mCherry reference marker across 12 independent experiments (N = 650 cells). Rpn11p-mCherry was consistently used for reference and the nuclear region is identified using GMM. Only time points with ≥25 dividing cells were included in the analysis. (**A**-**C**) Correlation of nuclear area with RLS. Pearson correlation (**A**), the corresponding -$\log_{10}$ p-values (**B**) , and the number of actively dividing cells (**C**) as a function of time are shown. (**D**-**F**) Correlation of cell area with replicative lifespan. Pearson correlation (**D**), the corresponding -$\log_{10}$ p-values (**E**) , and the number of actively dividing cells (**F**) as a function of time are shown. (**G**-**I**) Correlation of nuclear-to-cell area ratio with RLS. Pearson correlation (**G**), the corresponding -$\log_{10}$ p-values (**H**), and the number of actively dividing cells (I) as a function of time are shown. Blue dots indicate times at which the -log10(P) reaches the maximum.

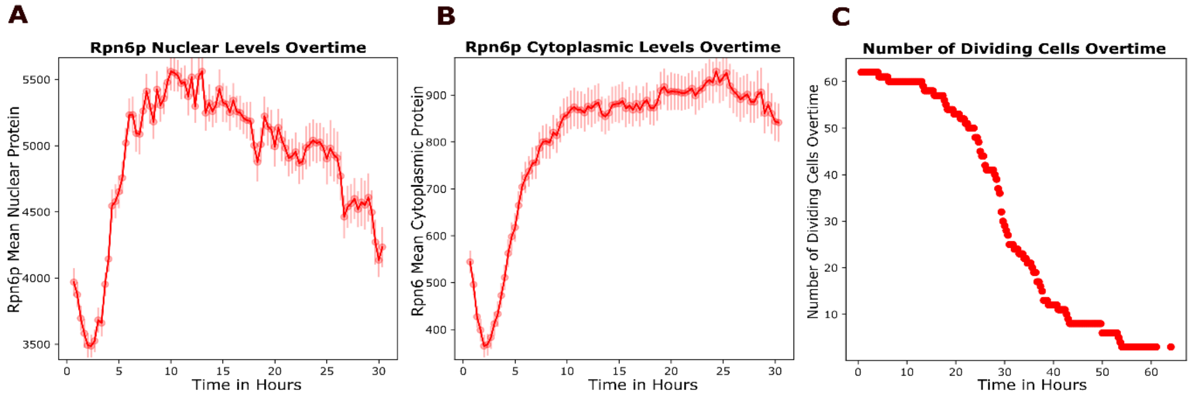


**Supplementary Fig. 5. Rpn6p reporter displays similar dynamics as Rpn11p** (**A**–**C**) Quantification of Rpn6p-mCherry localization over time in 62 dividing cells. Shown are nuclear Rpn6p intensity (**A**), cytoplasmic Rpn6p intensity (**B**), and number of dividing cells over time (**C**). Shaded error bars indicate the standard error of the mean (SEM). Nuclear and cytoplasmic intensities were extracted from single-cell tracking of Rpn6p-mCherry-expressing cells.

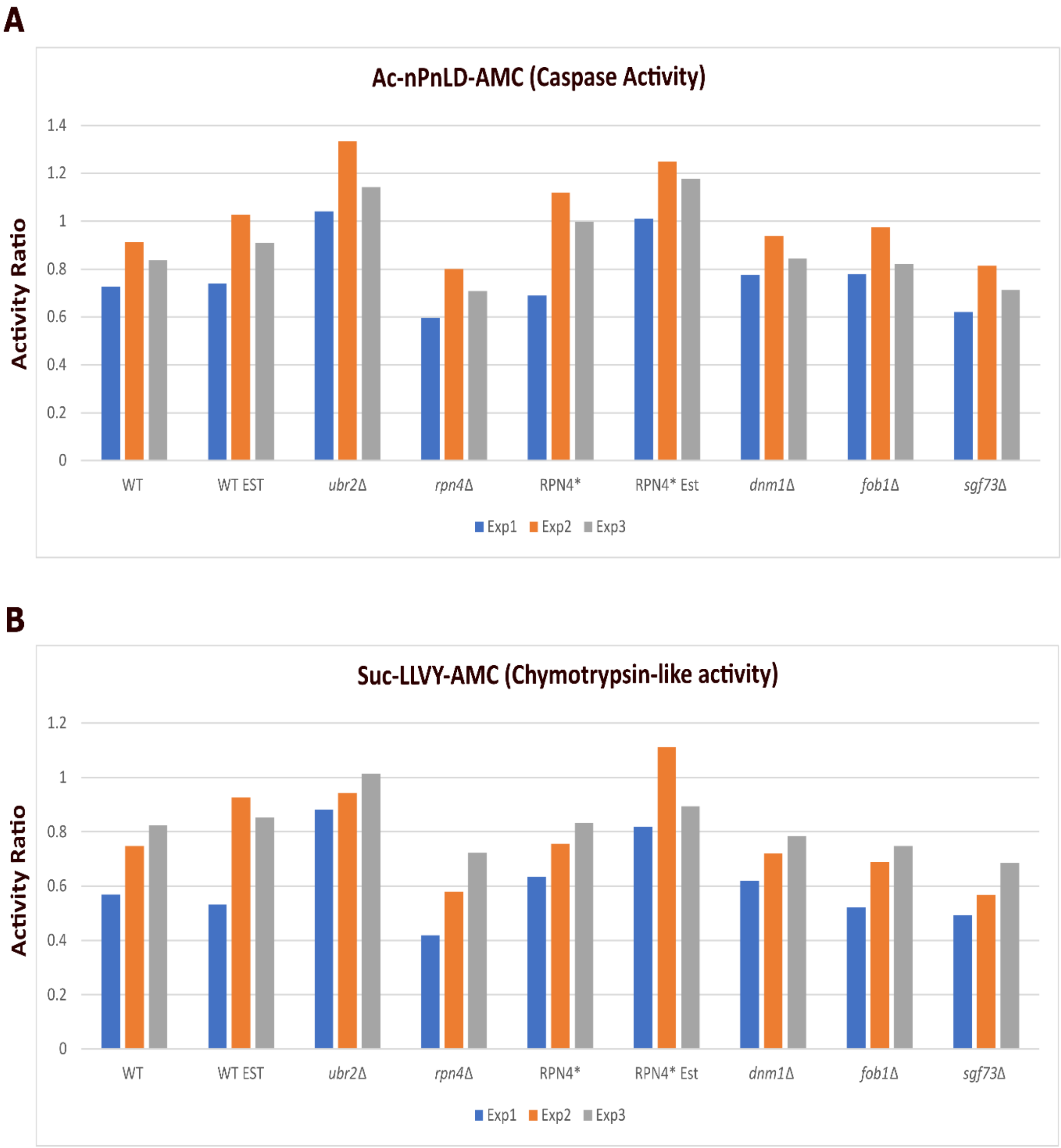


**Supplementary Fig. 6. Proteasome activity measured for different strains. (A)** Caspase-like proteasomal activity measured using the fluorogenic substrate Ac-nLPnLD-AMC. **(B)** Chymotrypsin-like proteasomal activity measured using the fluorogenic substrate Suc-LLVY-AMC. Strains tested include Wildtype (WT), WT treated with estradiol (WT EST), *ubr2Δ, rpn4Δ,* Rpn4*, Rpn4* treated with estradiol (RPN4* EST), *dnm1Δ*, *fob1Δ*, and *sfg73Δ*. For estradiol-treated samples, 16 nM estradiol was added for ~14 hours prior to harvesting. Cells were collected at an $OD_{600}$ of 0.8–1.0, and 50 µg of total protein was assayed in the presence of 100 µM fluorogenic substrate. Proteasomal activity was measured in three independent experiments (Exp1–Exp3) and is presented as the activity ratio, calculated by comparing reactions with and without 50 µg/ml of the proteasome inhibitor MG132 to confirm specificity.

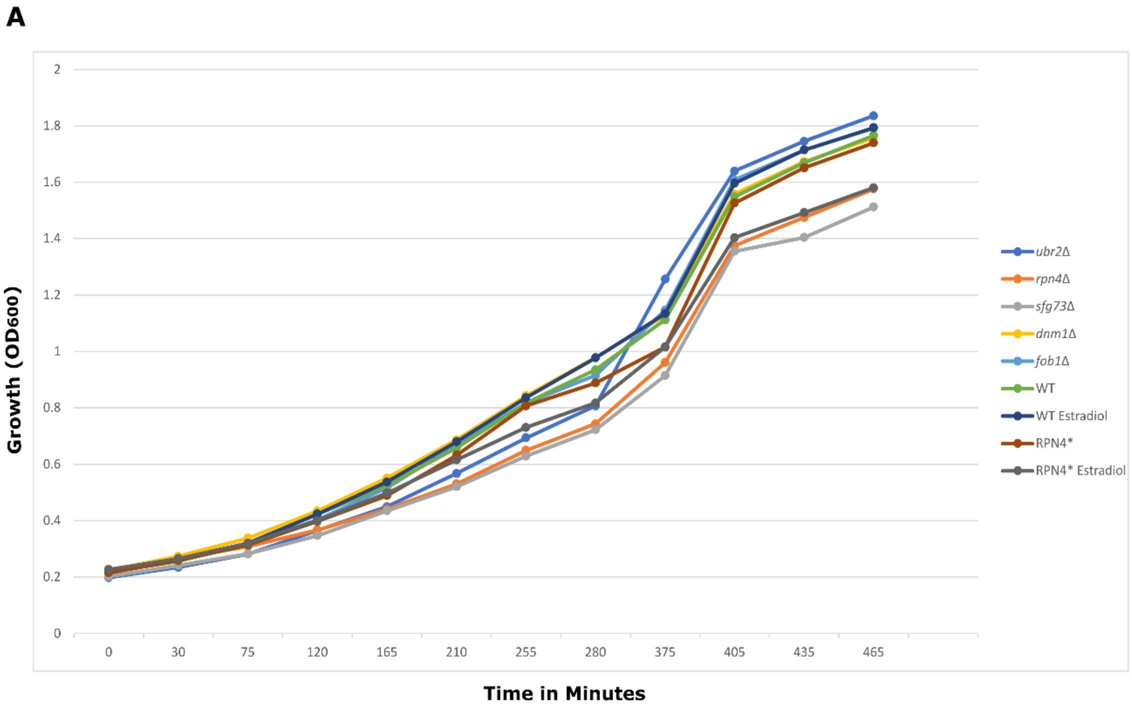


**Supplementary Fig. 7. Growth curves for different strains.** Growth measurements of *ubr2*Δ, *rpn4*Δ, *sfg73*Δ, *dnm1*Δ, *fob1*Δ, WT, WT estradiol, Rpn4*, and Rpn4* estradiol. Overnight cultures were diluted to an $OD_{600}$ of ~0.2 in galactose rich media and grown near stationary phase. Cell density ($OD_{600}$) was measured at intervals as indicated and used to plot the growth curves. Samples given 16nM of estradiol were incubated during the start of the experiment.

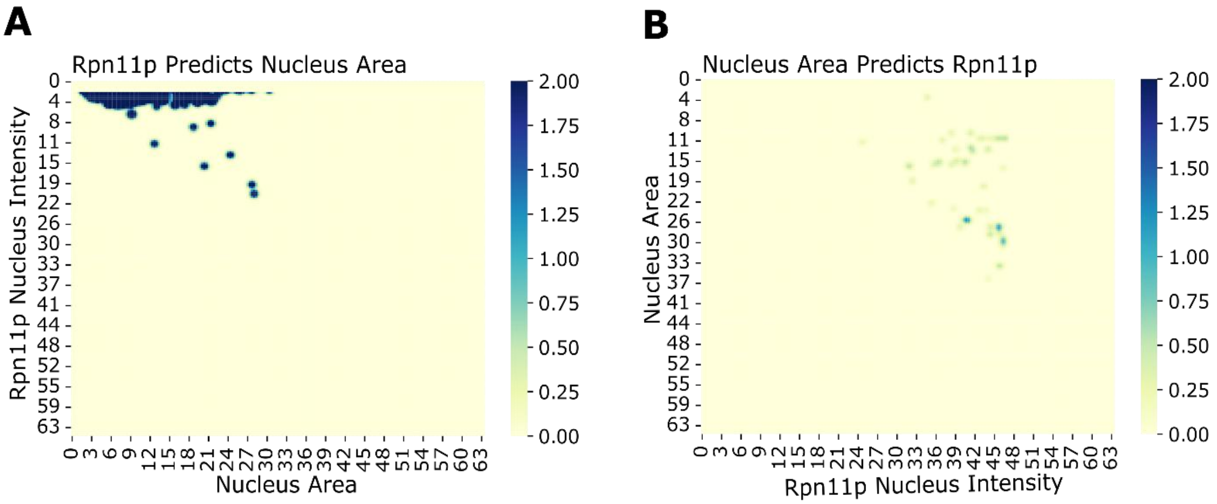


**Supplementary Fig. 8. Predicting temporal order using Granger causality with decision trees**. The following heatmaps display the -log10 P values calculated from an F-test. A Gaussian blur and threshold is used to filter background noise. (**A)** using Rpn11p nuclear intensity to predict nucleus area in later times. (**B**) using nuclear area to predict Rpn11p nucleus intensity in later times. The time points for both the y-axis and x-axis are the time in hours. Dataset is an accumulation of our Rpn11p-mCherry strains with ~650 cells. Time points with less than 56 dividing cells are excluded. Color scale indicates -log10(P) values.

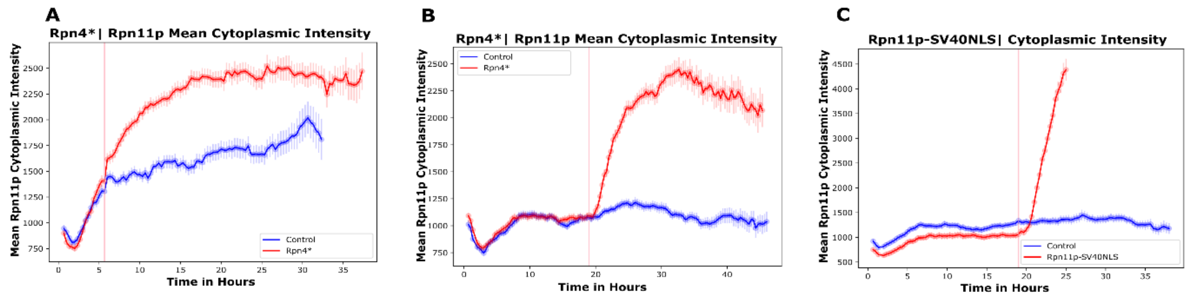


**Supplementary Fig. 9. Dynamics of cytoplasmic Rpn11p intensity following Rpn4* and Rpn11-SV40NLS induction.** (**A**) Time-course of mean cytoplasmic fluorescence intensity of Rpn11p in control cells (blue) versus cells expressing truncated Rpn4* (red) induced at 5.5 h (indicated by the red vertical line). (**B**) As in (**A**), but with induction at 19.2 h. (**C**) Time-course of mean cytoplasmic fluorescence intensity of Rpn11p in control cells (blue) versus cells being induced with Rpn11p–SV40NLS (red) at 19.2 h (red vertical line). Shaded areas denote SEM.

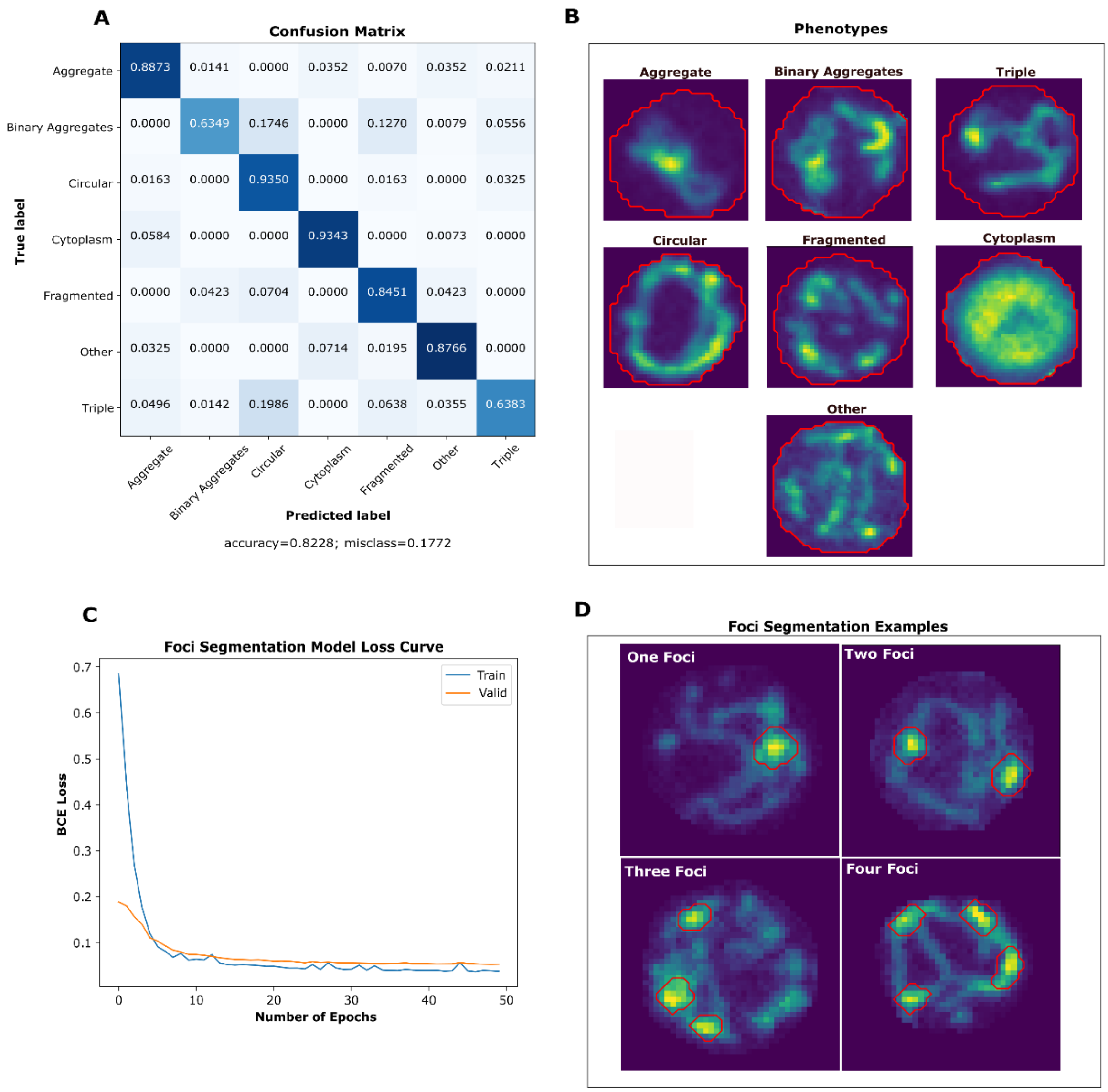


**Supplementary Fig. 10. Classification of mitochondrial morphologies**. (**A**) A confusion matrix of the ResNet50 model predictions for the validation dataset, with true labels on the y-axis and predicted labels on the x-axis. The validation dataset encompasses diverse experiments, integrating different protein organelle tags and knockout experiments. (**B**) Representative images of identified phenotypes. The red circle delineates the cell boarder. (**C**-**D**) relate to the Foci Segmentation Model. (**C**) the loss curve, with the training data depicted by the blue curve and the validation data by the orange curve. The y-axis signifies the Binary Cross-Entropy with Logits Loss (BCE), while the x-axis denotes the number of epochs. (**D** a selection of mitochondrial phenotypes with foci highlighted in red. The foci count per cell is outlined in white, ranging from one to four. The model can detect up to four or more foci.

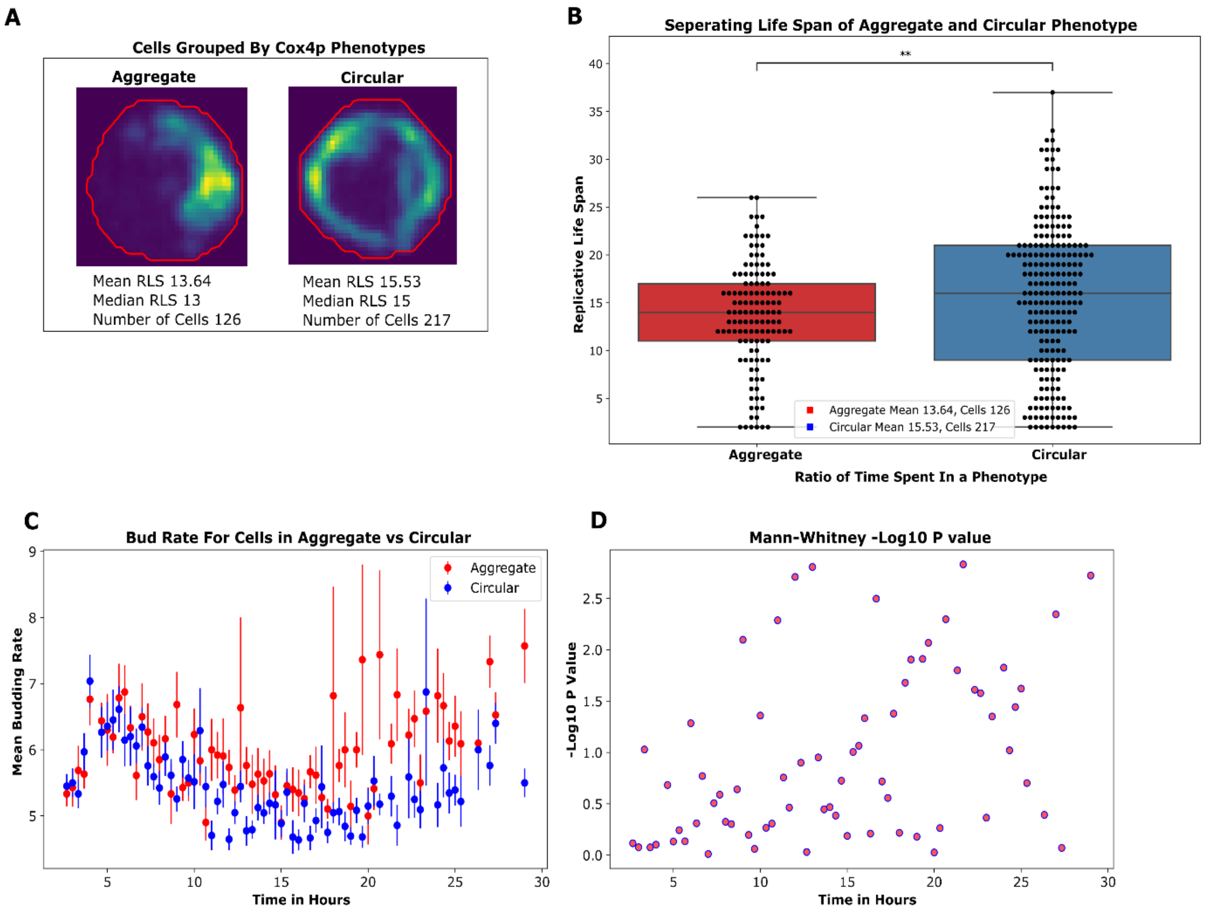


**Supplementary Fig. 11. Cells with circular mitochondrial phenotype live longer and bud slower than those with aggregate phenotype.** (**A**) example images of the aggregate and circular phenotype. For simplicity, cells displaying binary and triple phenotype are reclassified into the aggregate state. Cells displaying the fragmented state are recalssified into the circular state. This is decided by observing phenotype similarities to the aggregate or circular state. Next cells are grouped into aggregate or circular based on the time spent in the designated state. (**B**) a boxplot of the RLS for each cell classified in the aggregate or circular state. P value based on Mann-Whitney test is 9.605e-03. (**C**) the mean budding rate of cells with circular and aggregate phenotyes over time. Times with less than 10 dividing mother cells are excluded. The SEM of cells dividing at that time is depicted as error bars. (**D**) -log10 P values calculated using a Mann-whitney U test comparing the budding rate of cells calssified in the aggregate or circular state. The dataset consolidates our Cox4p-GFP and Rpn11p-mCherry strains from six independent experiments, encompassing 343 cells.

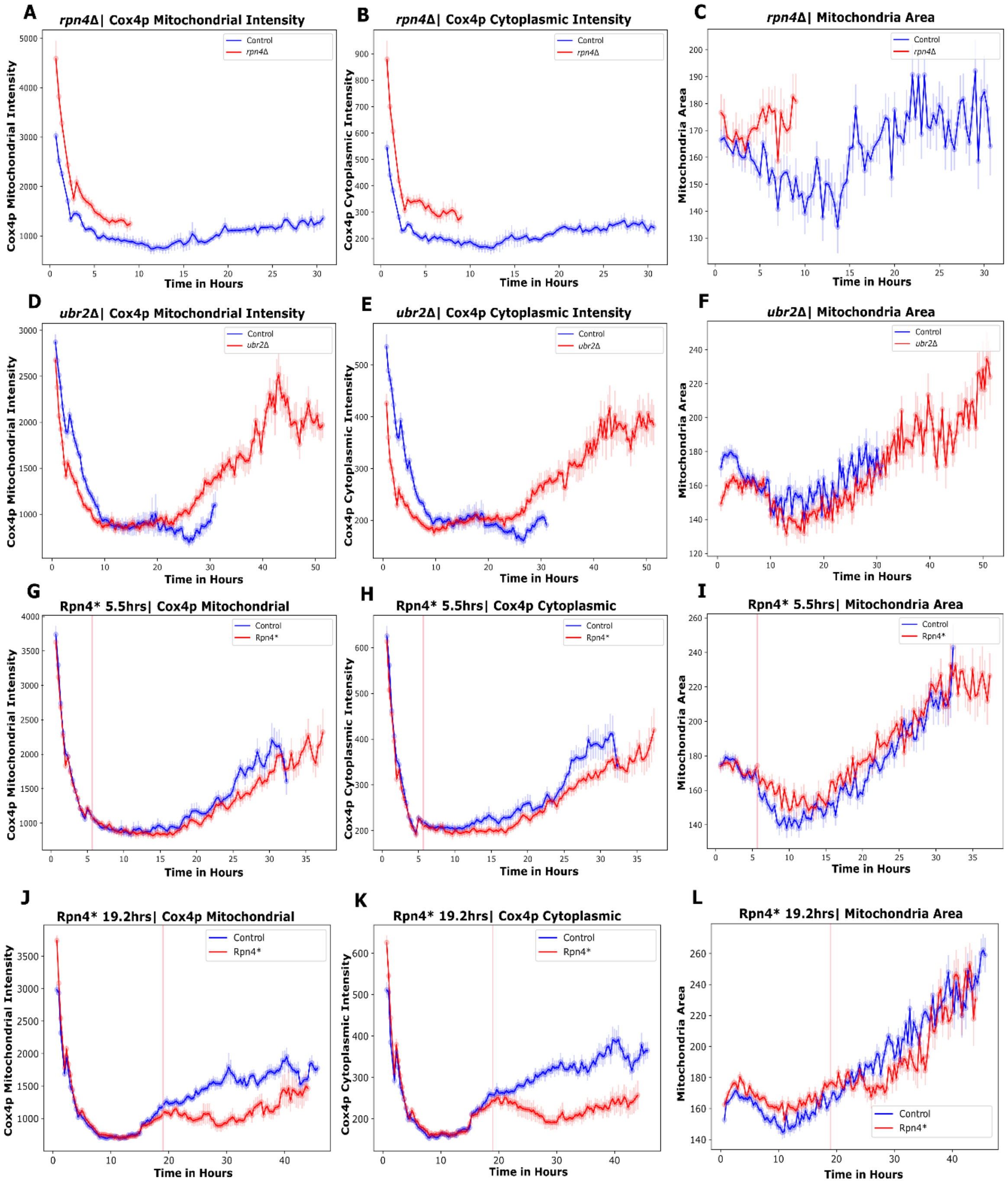


**Supplementary Fig. 12. Response of Cox4p signal to various proteasome perturbations.** The mean Cox4p mitochondrial intensity, mean Cox4p cytoplasmic intensity, and mean inner mitochondrial area of living dividing cells are shown for different proteasome perturbations of *rpn4*Δ (**A**-**C**), *ubr2*Δ (**D**-**F**), and temporal induction of Rpn4* at time 5.5hrs (**G**-**I**), and 19.2hrs (**J**-**L**). Only time points with a minimum of 15 cells are displayed. The blue curves represent the control, and the red curves represent the proteasome perturbation. SEM is displayed in the shaded areas within the curves. The pink line indicates when induction occurs. Both Rpn4* experiments were performed twice, and the data represents the accumulation of the separate experiments. Number of cells: For **A**-**C**, N= 47 for *rpn4*Δ and N=47 for the control. For **D**-**F**, N=61 for *ubr2*Δ and N= 46 for the control. For **G**-**I**, N=103 for

Rpn4* induction at 5.5 hours and N=107 for the control. For **J-L**, N= 115 for Rpn4* induction at time 19.2 hours and N=114 for the control.

**A**

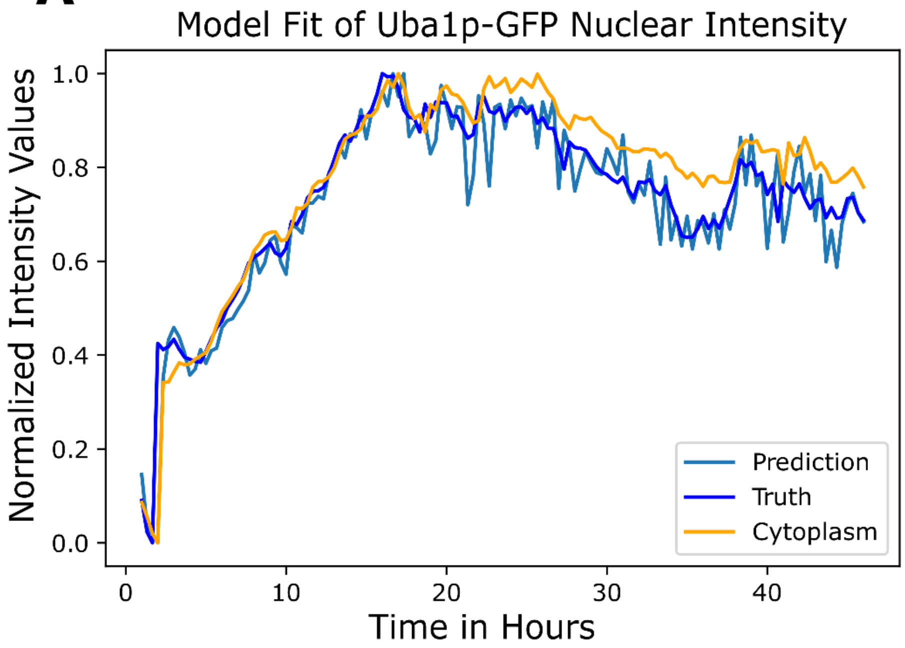


**Supplementary Fig. 13. Dynamics of Uba1p intensity in nucleus (dark blue) and cytoplasm (orange).** The light blue curve is the fit by the model,

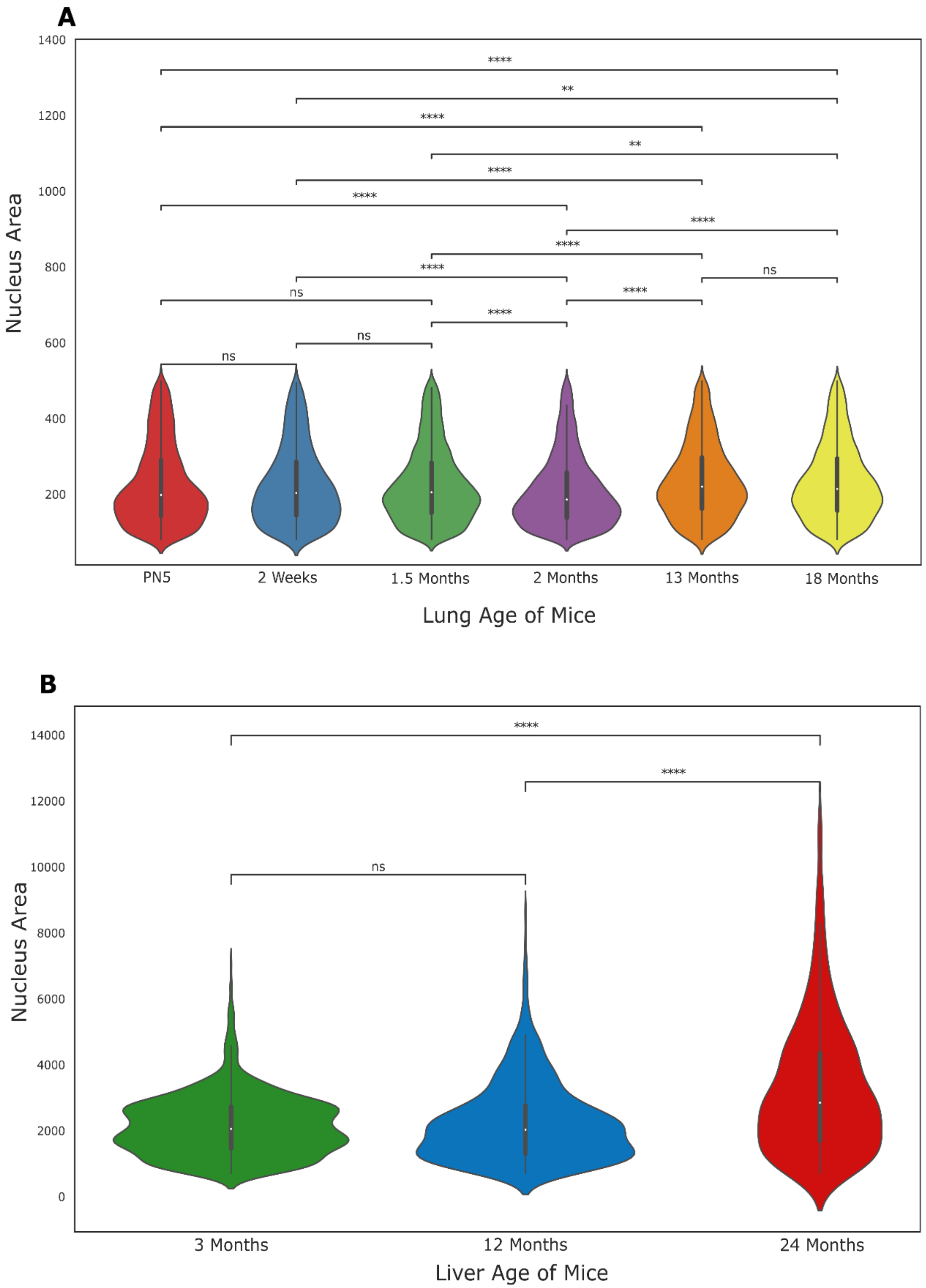


**Supplementary Fig. 14. The nucleus area of old mouse lung and liver cells are larger than those in young mice.** (**A**) Violin plot of nucleus area in lung tissue across different ages. Postnatal day 5 (PN5) and mice aged 2

weeks, 1.5 months, 2 months, 13 months, and 18 months are shown. The y-axis represents nucleus area, and the white dot denotes the median. The number of nuclei analyzed per group: PN5 (n = 6,016), 2 weeks (n = 2,568), 1.5 months (n = 13,041), 2 months (n = 9,288), 13 months (n = 3,385), and 18 months (n = 3,900). Statistical significance was determined using the Mann-Whitney U test, with $p < 0.05$ indicated by "*". (**B**) Violin plot of nucleus area in liver tissue across different ages. Mice aged 3, 12, and 24 months are shown. The y-axis represents nucleus area, and the white dot denotes the median. The number of nuclei analyzed per group: 3 months (n ≈ 1,034), 12 months (n = 693), and 24 months (n = 616). Statistical significance was determined using the Mann-Whitney U test, with $p < 0.05$ indicated by "*". The p-value for 12 vs. 24 months = $9.93 \times 10^{-22}$, and for 3 vs. 24 months = $8.25 \times 10^{-29}$.
