## Supplemental Table 2 for "Yeast aging from a dynamic systems perspective: Analysis of single cell trajectories reveals significant interplay between nuclear size scaling, proteasome dynamics, and mitochondrial morphology"

### Key:

- $S_n$ – Surface area of the nucleus at (t-1)
- $\Delta S_n$ – Surface area difference between (t-1) and (t)
- $C_0$ – amount of transport per unit time and per unit area and per unit of cytoplasmic protein concentration
- $C_1$ – Density and speed of the transporter
- $v_n$ – Volume of nucleus (t-1)
- $\Delta v_n$– Change in Volume of nucleus between two time points (t-1) and (t)
- $I_C$ – Cytoplasmic protein intensity at (t-1)
- $\Delta t$ – Difference between two time points
- $I_n$ – Nuclear protein concentration (t-1)
- $\Delta I_n$ – Change in nuclear protein concentration between (t-1) and (t)
- $\gamma$ – protein degradation rate
- $C_0 S_n \Delta t$ – represents the protein flux within the nucleus
- $\Delta(v_n I_n)$ – the total change between the nuclear volume and nuclear protein concentration

### Assuming Spherical Shape:

$$S_n = 4\pi R^2\,,\ \frac{s_n}{v_n} = \frac{3\sqrt{4\pi}}{\sqrt{s_n}}\ ,\ C_1 = C_0 \cdot 3\sqrt{4\pi}$$

$$v_n = \frac{4\pi}{3}R^3,\ \frac{\Delta v_n}{v_n} = \frac{3}{2}\frac{\Delta s_n}{s_n}$$

### Equation Derivation:

1. $I_C \cdot\ C_0 S_n \Delta t =\ \Delta(v_n I_n) + \gamma \cdot I_n v_n \Delta t$

2. $I_C \cdot\ C_0 S_n \Delta t =\ \Delta v_n I_n + v_n \Delta I_n + \gamma I_n v_n \Delta t$

3. $\Delta I_n = I_C C_0 \frac{s_n}{v_n} \Delta t -\ I_n \left(\frac{\Delta v_n}{v_n}\ + \gamma \Delta t\ \right)$

4. $\Delta I_n = \frac{C_1 I_C \Delta t}{\sqrt{s_n}} -\ I_n( \frac{3}{2}\frac{\Delta s_n}{s_n}\ + \gamma \Delta t\ )$
