## Supplemental Table 3 for "Yeast aging from a dynamic systems perspective: Analysis of single cell trajectories reveals significant interplay between nuclear size scaling, proteasome dynamics, and mitochondrial morphology"

| | C1 | Gamma |
|---|---|---|
| Long Lived | 40.4 | 0.8481 |
| Short Lived | 40.4 | 0.8481 |
| Wildtype | 40.4 | 0.8481 |
| Single Cell | 40.4 | 0.8481 |
| Initial | 42.2 | 0.9042 |
| 5.5h | 25.8 | 0.66 |
| 19.2hrs | 26 | 0.663 |
| 5.5hr Control | 30.8 | 0.66 |
| 19.2hrs Control | 30.2 | 0.66 |
| Uba1p | 24.8 | 0.85 |
